# Entorhinal velocity signals reflect environmental geometry

**DOI:** 10.1101/671222

**Authors:** Robert G K Munn, Caitlin S Mallory, Kiah Hardcastle, Dane M Chetkovich, Lisa M Giocomo

## Abstract

The entorhinal cortex contains neural signals for representing self-location, including grid cells that fire in periodic locations and velocity signals that encode an animal’s speed and head direction. Recent work revealed that the size and shape of the environment influences grid patterns. Whether entorhinal velocity signals are equally influenced or provide a universal metric for self-motion across environments remains unknown. Here, we report that changes to the size and shape of the environment result in re-scaling in entorhinal speed codes. Moreover, head direction cells re-organize in an experience-dependent manner to align with the axis of environmental change. A knockout mouse model allows a dissociation of the coordination between cell types, with grid and speed, but not head direction, cells responding in concert to environmental change. These results align with predictions of grid cell attractor models and point to inherent flexibility in the coding features of multiple functionally-defined entorhinal cell types.

## Introduction

Navigation is a complex cognitive process requiring the integration of multi-sensory cues to form a unified percept of an animal’s position in space. The neural substrates for generating this position estimate are thought to reside in the medial entorhinal cortex (MEC) and include MEC grid cells, which fire in multiple spatial locations arranged in a hexagonal lattice ^1,2^. The emergence of periodicity in grid cell firing patterns despite frequent changes in an animal’s running speed and direction led to the proposal that grid cells actively use self-motion cues to build a metric representation of the local spatial environment ^1,3^. This self-motion information may be derived from MEC velocity signals, including speed cells, which change their firing rate as a function of running speed and head direction cells, which fire maximally when an animal faces a particular direction ^4–7^. However, recent work has revealed that grid cells do not provide an invariant spatial metric across all environments and instead grid patterns deform, distort, and rescale in response to the geometric shape of local environments ^8–11^. Sensory landmark cues, such as environmental boundaries, play a key role in driving such structural changes to grid patterns ^12–14^. It remains unknown, however, whether velocity signals show flexibility in their coding in response to metric changes to the environment, or contribute to environmentally driven changes in grid patterns. For example, MEC speed cells have been proposed to invariantly code for running speed across spatial contexts but these signals have not been broadly considered under conditions in which the geometric size and shape of an environment is altered ^4^.

Attractor network models, a class of computational models capable of generating grid cell firing patterns, provide a principled framework for how velocity signals may respond to environmental perturbations ^15^. In attractor network models, grid cell firing results from the invariant translation of periodic activity bumps across a neuronal lattice ^3,15,16^. Inputs that reflect the running speed and direction of the animal drive the translation of the activity bumps and thus, determine the spatial scale and structure of the resulting grid firing patterns. Faster translation of the activity bumps, driven by stronger velocity inputs, results in smaller grid spatial scales ^3,15,17–19^. These models make specific, untested predictions regarding how velocity signals should change in response to environmental perturbations. For example, parametrically decreasing the size of an open arena from a square to a smaller rectangle asymmetrically re-scales the grid pattern, with the distance between grid firing nodes decreasing only along the re-scaled axis ^10,11,20^. Attractor models can replicate this asymmetric grid re-scaling by asymmetrically re-scaling the velocity input to the grid network ^15^. Whether this occurs experimentally however, remains unknown.

Here, we use single-cell in vivo electrophysiology to record from grid, head direction and speed cells in MEC as mice explore familiar and transiently compressed or expanded open arenas. We examine whether speed and head direction signals provide invariant self-motion signals across environments or change their coding in response to metric changes to the environment. To then more directly assess whether speed or head direction signals contribute to the re-scaling observed in grid cells, we take advantage of the altered MEC speed signals observed after the loss of HCN channels in mutant mice ^21^.

## Results

### Behavioral paradigm

To examine how the firing patterns of MEC cells change after local environmental perturbations in mice, we recorded neurons in the superficial layers of MEC as mice foraged for randomly scattered food rewards in open arenas (see Methods). Mice explored either a familiar square followed by a compressed rectangle (compression condition: mouse n = 25, cell n = 193) or a familiar rectangle followed by an expanded square (expansion condition: mouse n = 16, cell n = 282). Neuron matching between environments was determined by cluster matching and determination of cluster center of mass (Figure S1). We identified cells as encoding position (P), head direction (H), or running speed (S) in their firing pattern using a statistical model based approach ^6,22^ (see Methods). Quantifications of tuning curve features were then used for additional classification, such as identifying P-encoding grid and border cells ^23^.

### Grid spacing in mice distorts after local environmental perturbation

We first considered grid cell patterns in mice after local environmental perturbations (compression mouse n = 14, grid cell n = 31; expansion mouse n = 5, grid cell n = 28). We created a re-scaled map by iteratively stretching and translating (up to 10 cm in the vertical and horizontal axes) the rate maps observed in the rectangular arena (Figure 1a-b, S2). We then quantified the amount of stretching (λ stretch proportion) and translation needed to best correlate the baseline map with the re-scaled map (Figure 1c, see methods). Consistent with observations in rats ^10,11^, the grid pattern re-scaled in response to changes in the local geometry of the environment (Figure 1c). In the compression condition, grid cells re-scaled asymmetrically. Compared to baseline, the grid pattern was more elliptical in the modified environment, with grid spacing decreasing in the axis of compression but remaining unchanged in the static axis (Figure 1e-f). In the expansion condition, grid cells re-scaled symmetrically. Grid spacing was larger in both the expanded and the static axes (Figure 1h), and the ellipticity of the grid pattern did not change (Figure 1e,h). In both conditions the spatial phase of grid maps also translated, but this did not occur in a systematic direction (Figure 1g). As in rats, grid re-scaling was more pronounced for larger compared to smaller grid scales (Pearson correlation coefficient: compression r = 0.47, p = 0.008; expansion r = 0.40, p = 0.03) (Figure 1i,j) ^11^. Critically, the extent of rescaling was unrelated to running speed, average coverage (Figure S3), number of exposures (< 13 per animal, Figure S4A) or dorsal-ventral recording depth (Figure S4b). These data demonstrate that grid firing patterns re-scale in mice after systematic perturbations in the geometric shape of the spatial environment.

**Figure 1.**
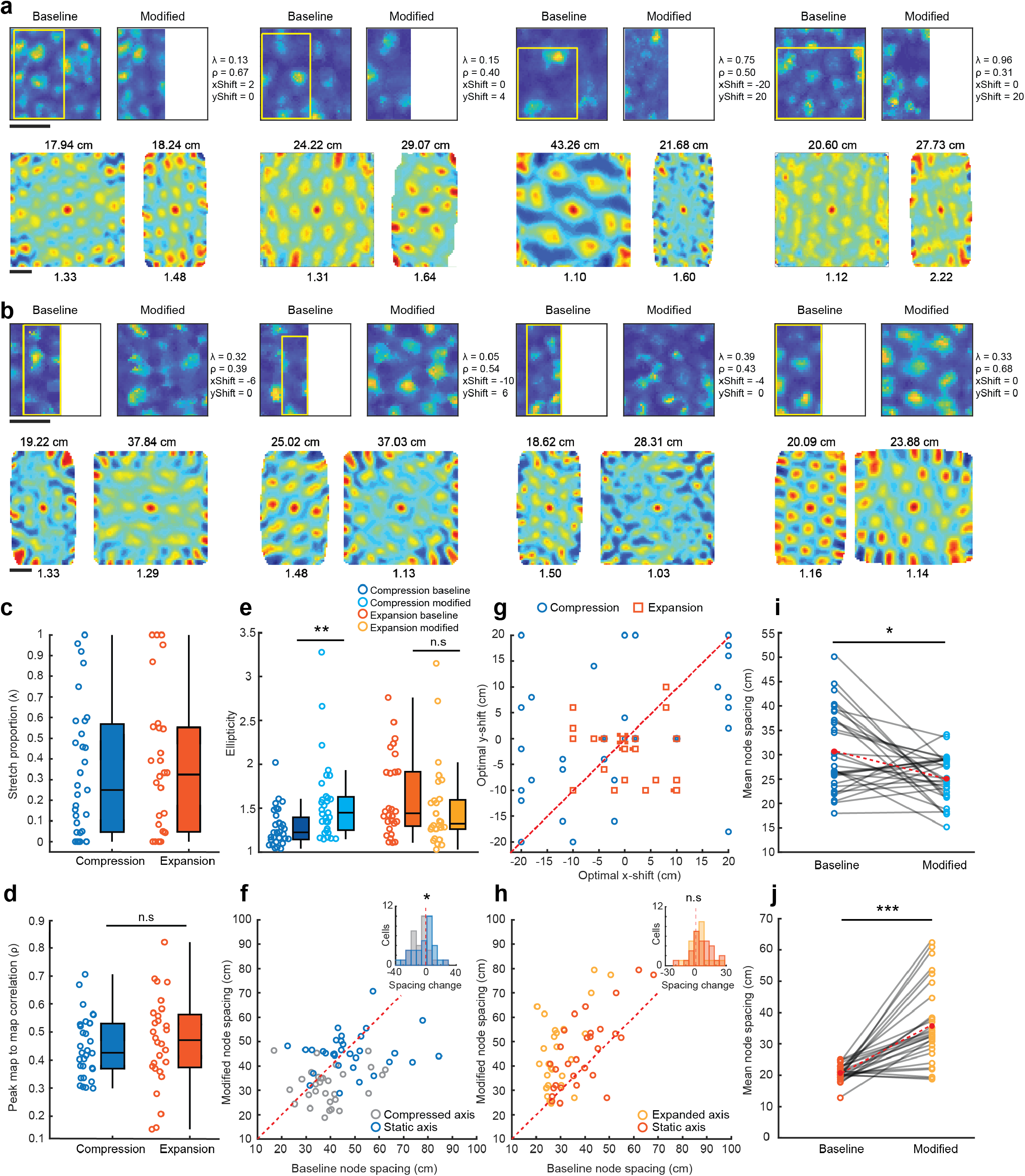
Environmental perturbation distorts grid spacing. **a,** Firing rate maps (top row) and spatial autocorrelations (bottom row) of grid cells in the baseline (left panels) and modified (right panels) environment for the compression condition. Maps are color coded for maximum (red) and minimum (blue) values. Scale bar indicates 50 cm. The area of the baseline environment that best correlated with the re-scaled map after stretching and translation is superimposed as a yellow rectangle. Stretch proportion (λ), magnitude of the peak correlation between baseline and rescaled maps (ρ) and optimal shift in X (xShift) and Y (yShift) at peak correlation are shown to the right of each rate map. Mean node spacing and ellipticity values are reported above and below the autocorrelation, respectively. **b,** Examples of grid cells labeled as in (a) for the expansion condition. **c,** Stretch proportion (λ) for grid cells in the compression (blue, n = 31) and expansion condition (orange, n= 28) (mean λ ± SEM; compression = 0.34 ± 0.057; expansion = 0.37 ± 0.065). Box illustrates median ± 25^th^ and 75^th^ percentiles and whiskers 1.5x the interquartile range. **d,** The magnitude of the peak correlation (ρ) between baseline and re-scaled maps is the same between compression (blue) and expansion (orange) conditions (mean correlation between baseline and rescaled map ± SEM; compression = 0.45 ± 0.02, expansion = 0.47 ± 0.03, Z = 0.77, p = 0.44). Data presented as in (c). **e,** Environmental compression caused an increase in grid ellipticity (mean ± SEM: compression baseline = 1.24 ± 0.25, modified = 1.59 ± 0.10, Z = 3.10, p = 0.002), while environmental expansion did not (expansion baseline = 1.62 ± 0.087, modified = 1.51 ± 0.091; Z = 1.18, p = 0.65). Data presented as in (c) **f,** In the compression condition, grid spacing decreased in the compressed but not the static axis (mean spacing ± SEM: compressed axis baseline = 38.97 ± 1.90 cm, modified = 32.56 ± 1.40 cm, Z = 2.78, p = 0.005; static axis baseline = 48.66 ± 2.48 cm, modified = 45.63 ± 1.49 cm, Z = 0.49, p = 0.62). Unity between conditions is illustrated by a dotted red line. Inset histogram: distribution of spacing changes in each axis (KS = 0.36, p = 0.03). **g,** Amount of shift (cm) in the X and Y direction needed to produce the optimal map-to-map correlation between baseline and re-scaled maps (mean shift ± SEM: compression, x-shift = −1.55 ± 2.71 cm, y-shift = 0.90 ± 2.26 cm, Z = 0.85, p = 0.39; expansion, x-shift = 0.43 ± 1.19 cm, y-shift = −2.29 ± 1.01 cm, Z = 1.48, p = 0.14). Overlapping data points denoted with a small square symbol. **h,** In the expansion condition, grid spacing increased in the compressed and static axis (mean spacing ± SEM: expanded axis baseline = 29.71 ± 1.44 cm, modified = 33.22 ± 1.92 cm, Z = 2.25, p = 0.024; static axis baseline = 39.26 ± 2.15 cm, modified = 45.44 ± 2.90 cm, Z = 2.85, p = 0.004). Inset histogram: distribution of spacing changes in each axis (KS = 0.26, p = 0.17). **I,j,** Mean grid spacing of nodes used to calculate ellpticity for cells in the baseline and modified compression (i) and expansion (j) condition. The overall mean is shown in red. Spacing decreased in compression (mean spacing (cm) ± SEM: compression baseline = 30.66 ± 1.48, compression modified = 25.24 ± 0.84, Z = 2.45, p = 0.014) and increased in expansion (mean spacing (cm) ± SEM: expansion baseline = 20.85 ± 0.47, expansion modified = 35.73 ± 2.40, Z = 4.42, p < 0.001). All panels: *p < 0.05, ** p < 0.01, ***p < 0.001, n.s., not significant.

### Speed coding and grid spacing re-scale in concert

We next sought to determine if MEC velocity signals respond to environmental perturbations. We began by considering MEC speed cells that increase their firing rate with running speed ^4^. We fit a linear function to the model-derived tuning curves of the firing rate by running speed (FR/RS) relationship in S-encoding cells (mouse n compression = 11, expansion = 8; S-encoding cell n, positive slope in baseline, compression = 21; expansion = 30). Indicating that environmental perturbations impact speed coding, the FR/RS slope and intercept increased after environmental compression and decreased after expansion (Figure 2a-d). We then examined whether there was axis specificity in the slope changes we observed after environmental perturbation. We considered the spike rate tuning curves of S-encoding cells relative to the animal’s running speed in the compressed/expanded versus static axis (S-encoding cell n, positive slope in compressed/expanded or static axis in baseline, compression = 26, expansion = 36) (Figure 2e). After the environment was compressed, the slope was greater in the compressed compared to static axis, matching the asymmetry in grid spacing observed in the same condition (mean slope Hz/(cm/s) ± SEM: compressed axis = 0.11 ± 0.02, static axis = 0.05 ± 0.01, Z = 3.33, p < 0.001) (Figure 2f). In contrast, there was no difference in slope between the expanded and static axes in the expansion, matching the lack of asymmetry in the grid pattern in the same condition (mean slope Hz/(cm/s) ± SEM: expanded axis = 0.05 ± 0.01, static axis = 0.03 ± 0.01, Z = 1.24, p = 0.21) (Figure 2g). The observation of analogous asymmetric changes in speed cell slope and grid re-scaling in the compression condition, coupled with analogous symmetric changes in speed cell slope and grid re-scaling in the expansion condition, is highly consistent with predictions of attractor models of grid cell formation ^15^. Moreover, these results reveal that speed cells are not invariant across all spatial contexts and instead, like grid cell firing patterns, show non-uniform changes in their coding features in response to changes in environmental geometry.

**Figure 2.**
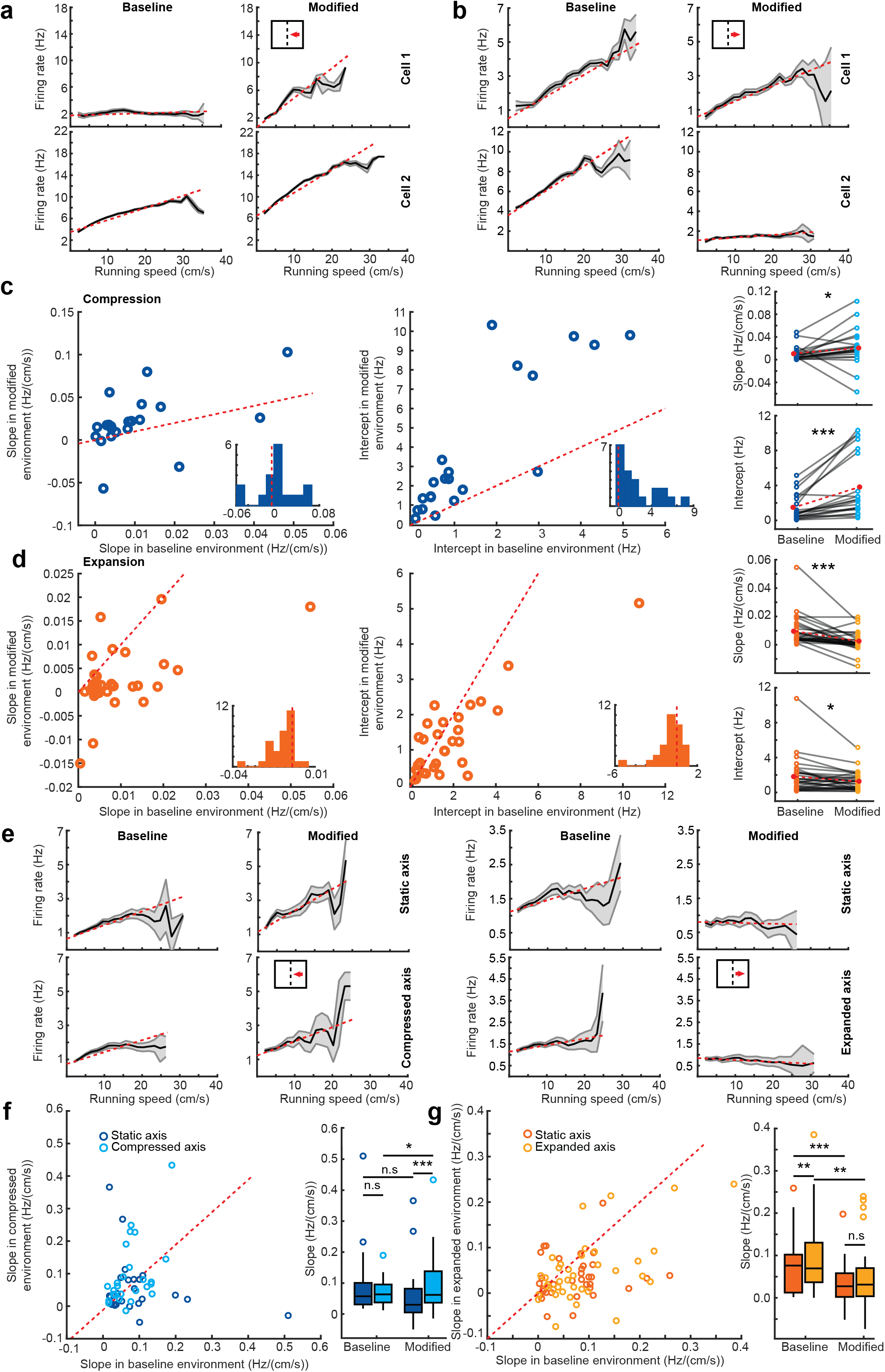
S-encoding cells re-scale in response to environmental perturbation. **a,** Two example S-encoding cells in the baseline (left panels) and modified (right panels) environment for the compression condition. Dotted red line indicates the linear function fit through the data. Solid black line indicates the mean response, with gray denoting ± SEM. **b,** Two example S-encoding cells labeled as in (a) for the expansion condition. **c,** Scatterplot of the slope (left) and intercept (middle) of S-encoding cells (n = 21) in the baseline and modified environment for the compression condition. The red dotted line illustrates unity between the baseline and modified environments. Inset: histogram of the difference in slope (left) or intercept (right) (modified – baseline). Red dotted line indicates zero. Far right panels show the change in response (slope, top; intercept, bottom) for each cell. The overall mean is shown in red (slope Hz/(cm/s) ± SEM: compression baseline = 0.011 ± 0.003, modified = 0.021 ± 0.007, Z = 2.14, p = 0.033; intercept (Hz) ± SEM: compression baseline = 1.50 ± 0.33, modified = 3.82 ± 0.78, Z = 3.84, p < 0.001) **d,** As in (c) for S-encoding cells recorded in the expansion condition (slope Hz/(cm/s) ± SEM expansion baseline = 0.010 ± 0.002, modified = 0.003 ± 0.001, Z = 4.00, p < 0.001; intercept (Hz) ± SEM: expansion baseline = 1.82 ± 0.37, modified = 1.28 ± 0.20, Z = 2.27, p = 0.023). **e,** Axis-specific firing rate by running speed responses for two example S-encoding cells, one in the compression (left) and one in the expansion (right) condition (static axis, top row; modified axis, bottom row). Labeled as in (a). **f,** Left: Slope of S-encoding cells in the static and compressed axis in the baseline versus modified environment. Asymmetry between axes is apparent in the modified but not baseline environment. Right: Boxplot illustrates median ± 25^th^ and 75^th^ percentiles and whiskers 1.5x the interquartile range. Single points beyond this range are plotted individually. **g,** As in (f) for cells recorded in the expansion condition. All panels: *p < 0.05, ** p < 0.01, ***p < 0.001, n.s., not significant.

Providing an additional readout of speed coding, EEG measured theta-band activity (5 – 11 Hz) in MEC increases in frequency as a function of running speed ^21,24^. Consistent with the increase in speed cell slope in the compression condition, the mean frequency of theta and intercept of theta frequency by running speed increased in the modified compared to baseline environment (n = 23 mice, n = 67 sessions, mean frequency (Hz) ± SEM: baseline = 8.66 ± 0.042, modified = 8.77 ± 0.039, Z = 4.12, p < 0.001; intercept (Hz) ± SEM: baseline = 8.33 ± 0.037, modified = 8.39 ± 0.041, Z = 2.89, p = 0.004) (Figure 3). Likewise, consistent with the decrease in speed cell slope in the expansion condition, the mean frequency of theta and intercept of theta frequency by running speed decreased in the modified compared to baseline environment (n = 11 mice, n = 74 sessions, mean frequency (Hz) ± SEM: baseline = 8.55 ± 0.024, modified = 8.49 ± 0.031, Z = 2.67, p = 0.007; intercept (Hz) ± SEM: baseline = 8.19 ± 0.036, modified = 8.13 ± 0.042, Z = 2.06, p = 0.039) (Figure 3). These results, together with our observations of nonuniform changes in speed cell coding, point to environmental perturbations modifying MEC speed signals at several levels of granularity.

**Figure 3.**
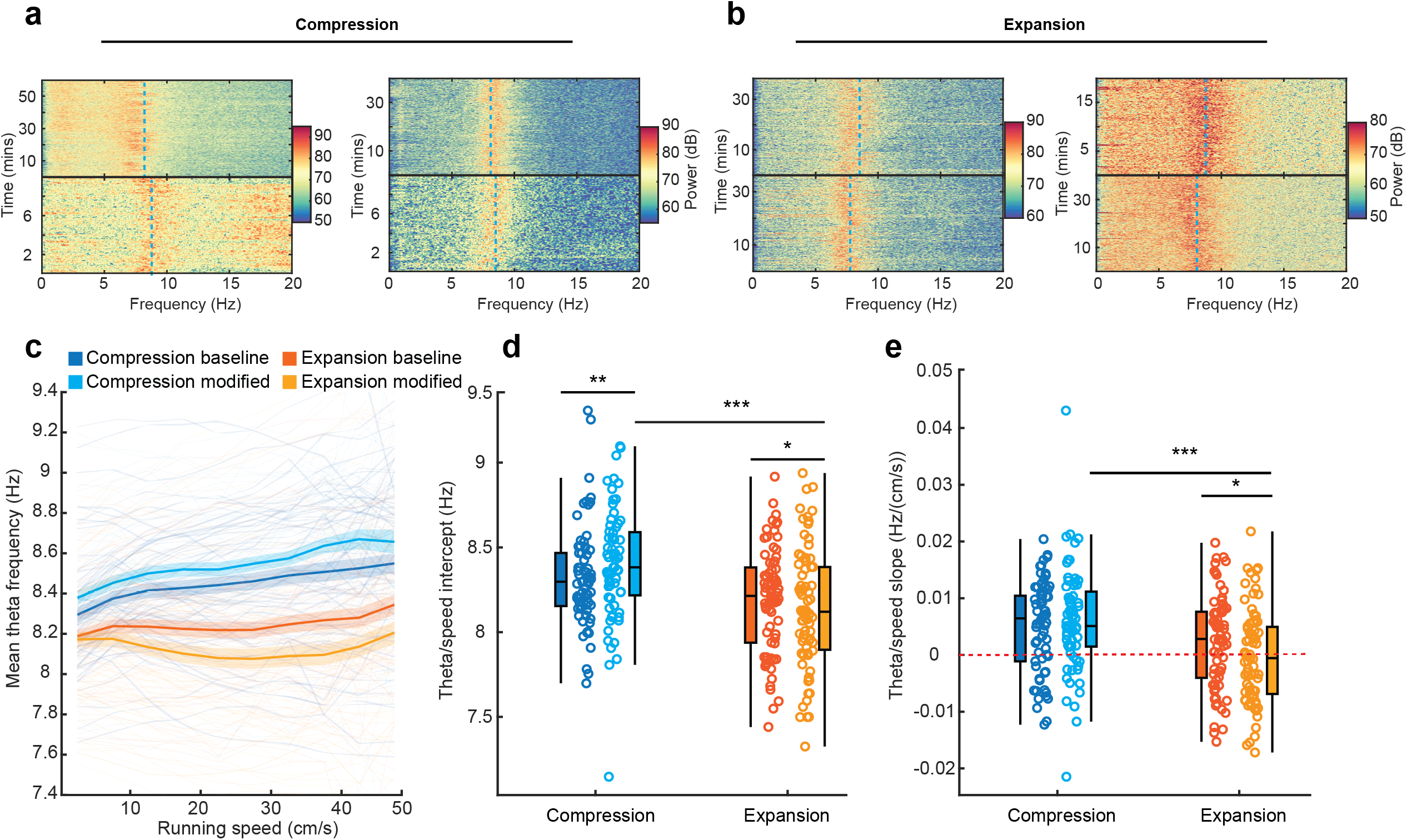
The gain of theta increases in compression and decreases in expansion conditions. **a,** Two example spectrograms showing LFP power in the 0-20 Hz frequency band for a recording session in the baseline (top panels) and modified (bottom panels) environments in the compression condition. A dotted blue line shows the mean theta frequency for that session. **b,** As in (a), but for two sessions (baseline, top and modified, bottom) in the expansion condition. **c,** Mean theta frequency by running speed for compression and expansion conditions. Solid lines show the mean, shaded region is ± SEM. Individual sessions are shown as faint lines. **d,** The intercept of linear fits through the frequency/speed relationship. **e,** The slope of linear fits through the theta frequency by running speed relationship. All panels: *p < 0.05, ** p < 0.01, ***p < 0.001, n.s., not significant.

### Experience-dependent asymmetry in head direction tuning

As a velocity signal consists of both speed and direction information, we next examined how head direction cells respond to environmental perturbations (compression mouse n = 17, H-encoding cell n = 49, expansion mouse n = 9, H-encoding cell n = 103). If head direction signals contribute to the asymmetric re-scaling of grid cells observed after environmental compression, a change in directional tuning should be observed primarily along the re-scaled axis. Interestingly however, we observed a bimodal bias in the preferred firing direction of H-encoding cells along the compressed/expanded axis only in the rectangular environment, regardless of whether this served as the modified environment for compression sessions or the baseline environment for expansion sessions (Hodges-Anje Omnibus test [H-A]: compression baseline m = 18, p = 0.53; modified m = 10, p < 0.001; expansion baseline m = 36, p < 0.046; modified m = 39, p = 0.19) (Figure 4a-b, Figure S5) ^25^. Moreover, the distribution of preferred phase angles differed between baseline and modified environments for both conditions, demonstrating a remapping of directional preference between baseline and modified environments (Watson-Williams test: compression F(1,96) = 9.33, p = 0.003; expansion F(1,204) = 11.53, p < 0.001). In the compression, this directional bias appeared attributable to many cells that aligned with the compressed axis in the baseline environment (preferred angle 45° – 135° and 225° – 315°) retaining their preferred direction in the modified environment (change in angle (°) ± SEM: baseline compressed axis aligned = 51.12 ± 9.83, baseline static axis aligned = 84.27 ± 10.27, Z = 2.55, p = 0.011) (Figure 4b-e). While many H-encoding cells changed their preferred directions, this did not occur with any systematic directional bias (change in angle (°) ± SEM: baseline expanded axis aligned = 76.36 ± 5.54, baseline static axis aligned = 83.36 ± 9.45, Z = 0.69, p = 0.49) (Figure 4b-d).

**Figure 4.**
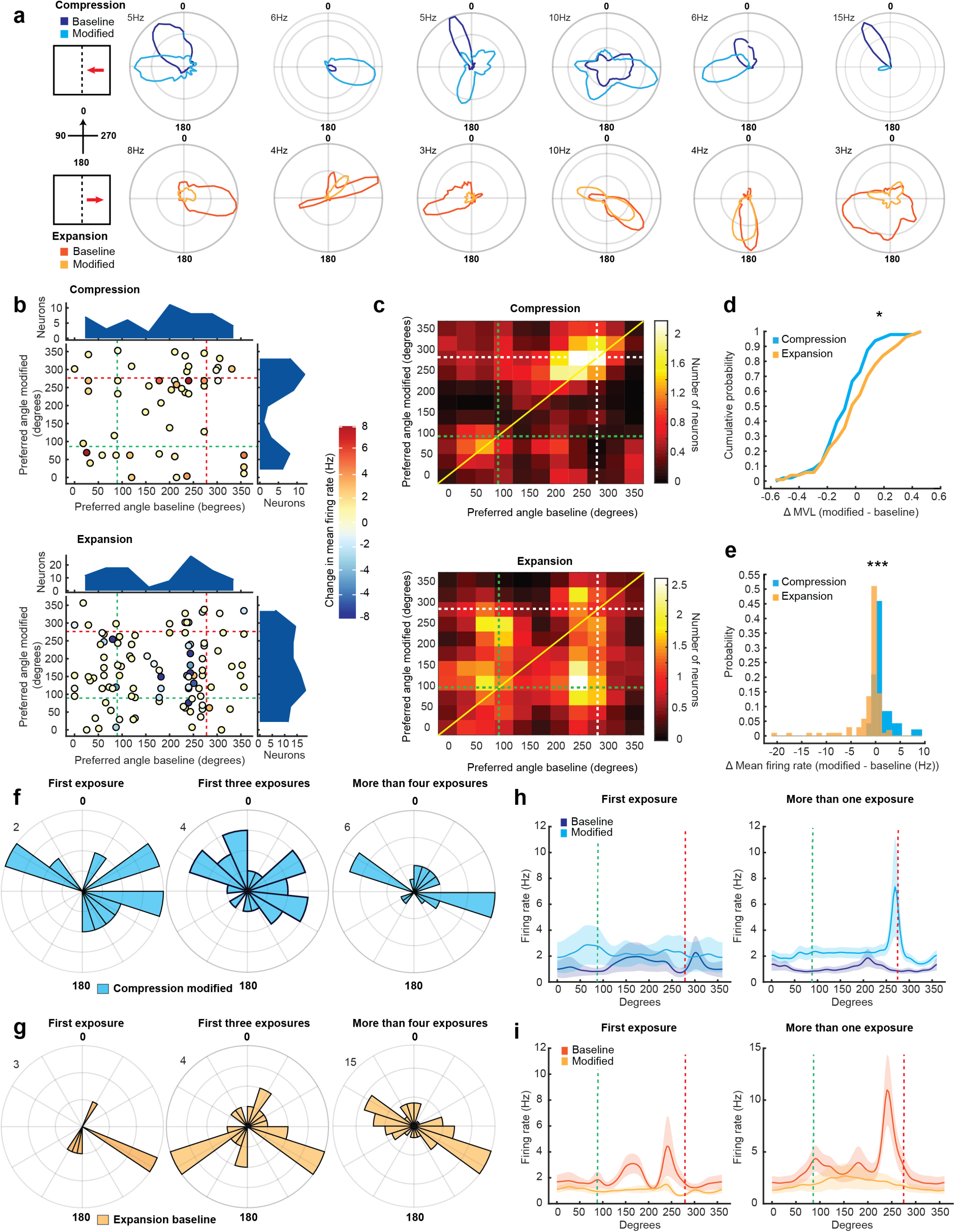
A directionally specific asymmetric bias develops after multiple exposures to modified environments. **a,** Polar plots of H-encoding cells in the compression (top row) and expansion (bottom row) condition. Top left numbers show the firing rate indicated by the outermost ring. **b,** Scatterplot of preferred phase angle (degrees) of H-encoding cells between baseline and modified environments in compression (top) and expansion (bottom) conditions. The change in firing rate between baseline and modified environments is indicated by colors coded for minimum (blue) and maximum (red) values. Marginal density plots (blue) indicate the distribution of phase angles. The wall axes are indicated by dotted lines (green, static wall; red, moved wall). **c,** The number of neurons falling in each 33° directional bin in the baseline and modified environments in the compression (top) and expansion (bottom) condition. A 2-bin Gaussian filter is applied to the data. The wall axes are indicated by dotted lines (green, static wall; white, moved wall). A solid yellow line demarks the bins in which there was no change in direction. **d,** The probability distribution of change in mean vector length (MVL) between environments for the compression (blue) and expansion (orange) condition (mean change in MVL ± SEM: compression = −0.06 ± 0.025, expansion = 0.013 ± 0.021, Z = 2.21, p = 0.027) **e,** The probability distribution of the change in mean firing rate between environments for the compression (blue) and expansion (orange) condition (Firing rate change (Hz) ± SEM: compression = 1.22 ± 0.29, expansion = −1.41 ± 0.35, Z = 7.07, p < 0.001). **f,g,** A bidirectional bias develops in H-encoding cells after multiple exposures to the modified environments. Polar histograms show the number of cells with a given preferred phase angle, with scale (n) indicated by the number on the top left. In the compression experiments (blue bars, (f)) no phase angle bias is evident on the first exposure to the compressed environment (left panel, n = 12) or over the first three exposures (n = 24). Cells recorded on sessions after the fourth exposure (n = 25; right panel) cluster bimodally. (g) shows the same as (f) but for cells recorded in the expansion condition (expansion left panel n = 7, middle n = 23 right n = 80). **h,I,** The directionally specific unimodal bias in firing rate develops with experience. H and I show the mean firing rate for all H-encoding cells in the compression (h) and expansion (i) conditions on the first exposure (left panels) and subsequent exposures (right panels). Solid lines are mean firing rates, ± SEM is shown as a shaded region. The wall axes are indicated by dotted lines (green, static wall; red, moved wall). All panels: *p < 0.05, ** p < 0.01, ***p < 0.001, n.s., not significant.

Firing rates were greatest in the asymmetric, rectangular environment, regardless of whether this environment served as the baseline or modified environment. In the compression, there was an overall increase in firing rates between baseline (square) and modified (rectangular) environments (mean FR ± SEM: baseline = 1.06 ± 0.19 Hz, modified = 2.28 ± 0.38 Hz, Z = 4.82, p < 0.001). In the expansion, firing rates decreased between baseline (rectangular) and modified (square) environments (mean FR (Hz) ± SEM: baseline = 3.19 ± 0.81, modified = 1.77 ± 0.59, Z = 5.50, p < 0.001) (Figure 4e). Unlike the bidirectional changes in preferred direction, the changes in firing rate appeared to be largely unidirectional, and primarily explained by changes in the directional bins that aligned with the moveable wall (Figure 4h-i).

The development of the directional bias in H-encoding cells depended on experience. There was no significant bias in directional preference in the modified compression or baseline expansion condition during the first, or first three, environmental perturbation sessions (H-A test, first exposure: compression modified n = 12, m = 2, p = 0.26; expansion baseline n = 7, m = 0, p = 0.11; H-A test, first three exposures: compression modified n = 7, m = 6, p = 0.19; expansion baseline n = 23, m = 7, p = 0.53) (Figure 4f-g). Starting with the fourth exposure however, the bimodal bias appeared in the modified compression and baseline expansion condition (H-A test, compression modified, n = 25, m = 4, p = 0.013, expansion baseline, n = 80, m = 18, p < 0.001) (Figure 4f-g). By contrast, the change in firing rate between environments was present on the first exposure to the modified environment (mean FR (Hz) ± SEM: compression baseline = 1.19 ± 0.43, modified = 2.31 ± 1.01, p = 0.01; expansion baseline = 1.90 ± 0.36, modified = 1.00 ± 0.19, p = 0.03) and persisted on three and more exposures (mean FR (Hz) ± SEM: compression baseline = 1.08 ± 0.25, modified = 2.46 ± 0.55, Z = 3.38, p < 0.001; expansion baseline = 4.03 ± 1.09, modified = 2.00 ± 0.76, Z = 5.37, p < 0.001). This experience dependent development of directional bias in the head direction signal contrasts with grid re-scaling, which does not vary with session number (Figure S4). Combined, these data suggest a dissociation between speed and directional signals, and that MEC head direction cells may not serve as the directional component of a velocity signal. Rather, head direction signals may reflect a learned signal regarding the direction in which the environment will change, or reflect learned knowledge of asymmetry in environmental geometry. The possibility also remains that, as recently suggested ^26^, only a small subset of segregated head direction cells serve the role of providing a velocity signal to the MEC grid network.

### HCN-channel auxiliary subunit TRIP8b deletion reduces grid re-scaling

One challenge to examining how MEC velocity signals influence grid cells is the heterogeneous nature of speed coding in MEC, with running speed encoded by both putative excitatory and inhibitory cells, as well as in the frequency of EEG measured oscillatory activity ^4–6,24,27^. To more directly dissociate the contribution of speed and head direction to grid cell rescaling after environmental perturbations, we next took advantage of the altered speed signals previously observed after the loss of forebrain HCN channels ^21,28,29^. Here, we considered grid, head direction and speed cells in mice lacking tetratricopepteide repeat-containing Rab8b-interacting protein (TRIP8b), an accessory subunit necessary for the insertion of HCN channels into the post-synaptic membrane, the loss of which significantly attenuates the HCN conducted hyperpolarization-activated current ^30^.

First, we examined grid cells in TRIP8b knockout (KO) mice in the compression condition (mouse n = 9, grid cell n = 27). Compared to wildtype (WT) mice, grid cells in KO mice showed significantly less re-scaling (mean λ stretch proportion ± SEM: WT = 0.34 ± 0.057, KO = 0.21 ± 0.065, KS = 0.44, p = 0.0047, Figure 5a-b) and did not change in ellipticity between baseline and modified environments (mean ellipticity ± SEM: baseline = 1.46 ± 0.079, modified = 1.55 ± 0.11, Z = 0.79, p = 0.43) (Figure 5c). Moreover, unlike grid cells in WT mice, there was no difference in grid spacing in either the compressed or static axis between baseline and modified environments (mean spacing (cm) ± SEM: compressed axis baseline = = 50.32 ± 2.02, modified = 49.16 ± 2.16, Z = 0.29, p = 0.77; static axis baseline = 40.67 ± 2.13, modified = 36.79 ± 1.78 cm, Z = 1.66, p = 0.097) (Figure 5d-e). The difference in grid re-scaling between WT and KO mice did not reflect differences in baseline spatial stability, overall poorer peak correlations between baseline and rescaled rate maps or larger amounts of translation in the spatial phase of grid maps (mean spatial stability ± SEM: WT = 0.35 ± 0.037, KO = 0.28 ± 0.035, KS = 0.29, p = 0.145); (Figure 5f-g). Together, these results indicate that the loss of HCN channels via TRIP8b KO renders grid cells significantly less sensitive to environmental deformation.

**Figure 5.**
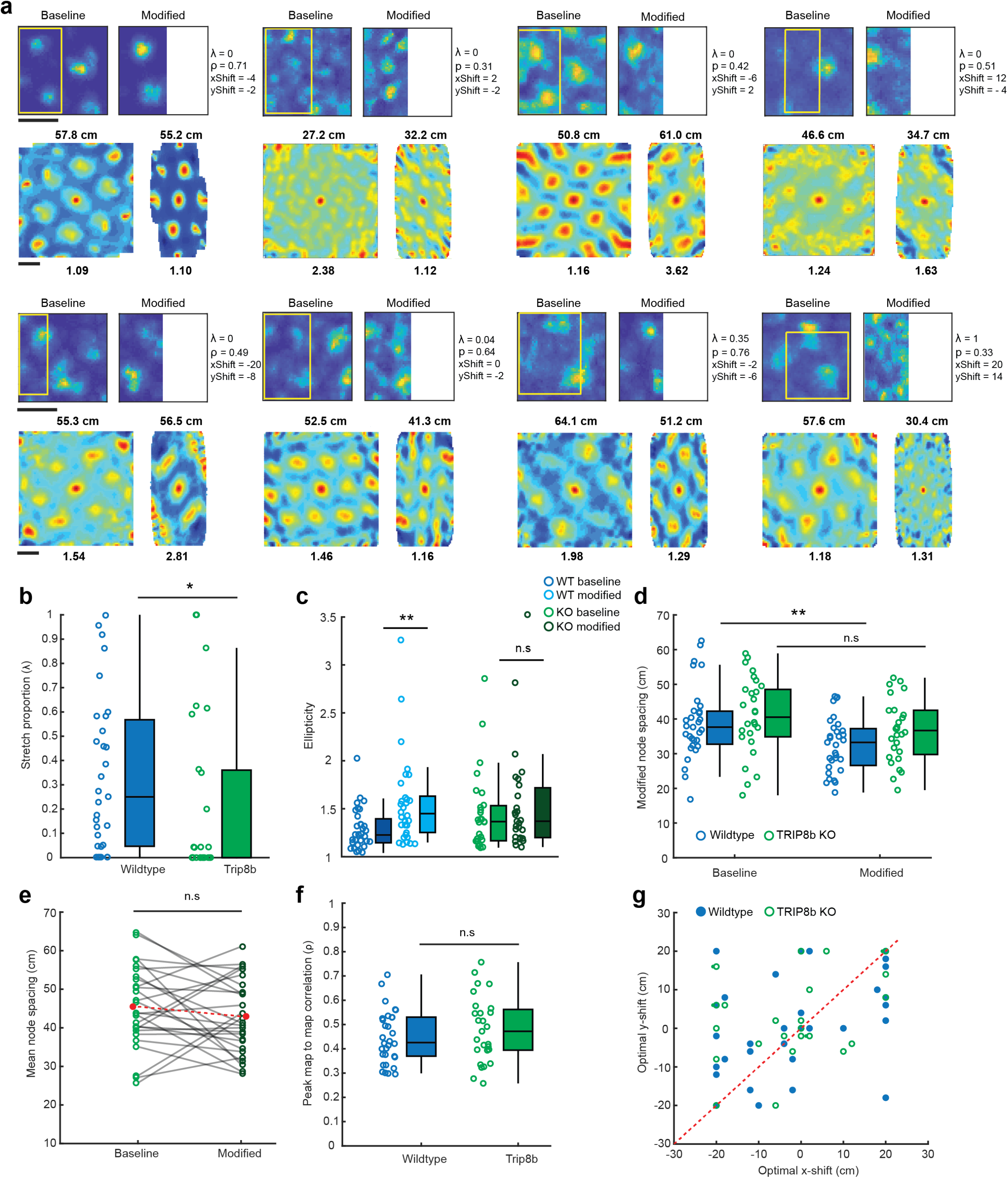
TRIP8b KO grid spacing is less sensitive to environmental perturbation. **a,** Firing rate maps (top rows) and spatial autocorrelations (bottom rows) of KO grid cells in the baseline (left panels) and modified (right panels) environment for the compression condition. Maps are color coded for maximum (red) and minimum (blue) values. Scale bar indicates 50 cm. The area of the baseline environment that best correlated with the re-scaled map after stretching and translation is superimposed as a yellow rectangle. Stretch proportion (λ), magnitude of the peak correlation between baseline and re-scaled maps (ρ) and optimal shift in X (xShift) and Y (yShift) at peak correlation are shown to the right of each rate map. Mean node spacing and ellipticity values are reported above and below the autocorrelation, respectively. **b,** Stretch proportion (λ) at optimum correlation. WT grid cells (blue) required more stretching than KO grid cells (green, n = 27) (mean A ± SEM: WT = 0.34 ± 0.06, KO = 0.21 ± 0.06, Z = 2.15, p = 0.031). Box illustrates median ± 25^th^ and 75^th^ percentiles and whiskers 1.5x the interquartile range. **c,** WT grid cells became significantly more elliptical compared to KO grid cells in the modified environment (Z = 2.50, p = 0.013). There was no difference in the ellipticity of KO grid cells between environments (mean ± SEM: KO baseline = 1.46 ± 0.08, modified = 1.55 ± 0.11, Z = 0.79, p = 0.43). Data are presented as in (b). **d,** In the compression condition, grid spacing in the compressed axis decreased in WT but not in KO mice (WT baseline = 39.0 ± 1.90cm, WT modified = 32.56 ± 1.40cm, Z = 2.78, p = 0.005; KO baseline = 40.67 ± 2.13cm, KO modified = 36.80 ± 1.78cm, Z = 1.66, p = 0.10). Data are presented as in (b). **e,** Mean grid spacing for cells in the baseline and modified environments for the compression condition. The overall mean is shown in red. There is no difference in spacing between environments (KO baseline = 45.50 ± 2.01 cm, KO modified = 42.97 ± 1.84 cm, Z = 1.20, p = 0.23). **f,** There was no difference in the peak map-to-map correlations between WT and KO grid cells (mean ± SEM: WT = 0.45 ± 0.02, KO = 0.50 ± 0.03, Z = 0.89, p = 0.37). Data are presented as in (b). **g,** There was no difference between WT and KO grid cells in the amount of shift in the X or Y direction needed to produce the optimal map-to-map correlation between baseline and re-scaled maps (mean ± SEM: x-shift, WT = −1.55 ± 2.71 cm, KO = −2.67 ± 2.67 cm, Z = 0.14, p = 0.89; y-shift, WT = 0.90 ± 2.26 cm, KO = 3.33 ± 2.16 cm, Z = 0.69, p = 0.49). Overlapping points denoted with a small square symbol. All panels: *p < 0.05, ** p < 0.01, ***p < 0.001, n.s., not significant.

### Speed coding remains stable after environmental perturbation in TRIP8b KO mice

To assess the relationship between speed coding and grid re-scaling after environmental perturbation, we examined the coding features of speed cells in TRIP8b KO animals in the compression condition (mouse n = 11, S-encoding cells n = 76) (Figure 6a). Similar proportions of cells encoded speed in WT and KO mice in baseline (WT = 27.46 %, n = 53/193, KO = 21.05 %, n = 16/76, X^2^ = 1.09, p = 0.30). However, compared to WT mice, very few KO S-encoding cells retained their speed coding in the compressed environment (WT = 12.04%, n = 23, KO = 3.94% n = 3, X^2^ =3.97 p = 0.046) (Figure 6b). Given this, we also considered speed coding using only a traditional score-based approach (speed score P95 = 0.045, speed cell n = 12; see Methods). However, even when considering this more broadly defined speed cell population, there was no difference in the FR/RS slope between baseline and modified environments in KO mice (mean slope (Hz/(cm/s)) ± SEM: baseline = 0.034 ± 0.007, modified = 0.027 ± 0.007, p = 0.73) (Figure 6c-d). This contrasted with a significant increase in slope in WT mice when using the same cell classification criterion (speed cell n = 58, mean slope (Hz/(cm/s)) ± SEM: baseline = 0.042 ± 0.006, modified = 0.057 ± 0.009, Z = 2.31, p = 0.02, Figure S5).

**Figure 6.**
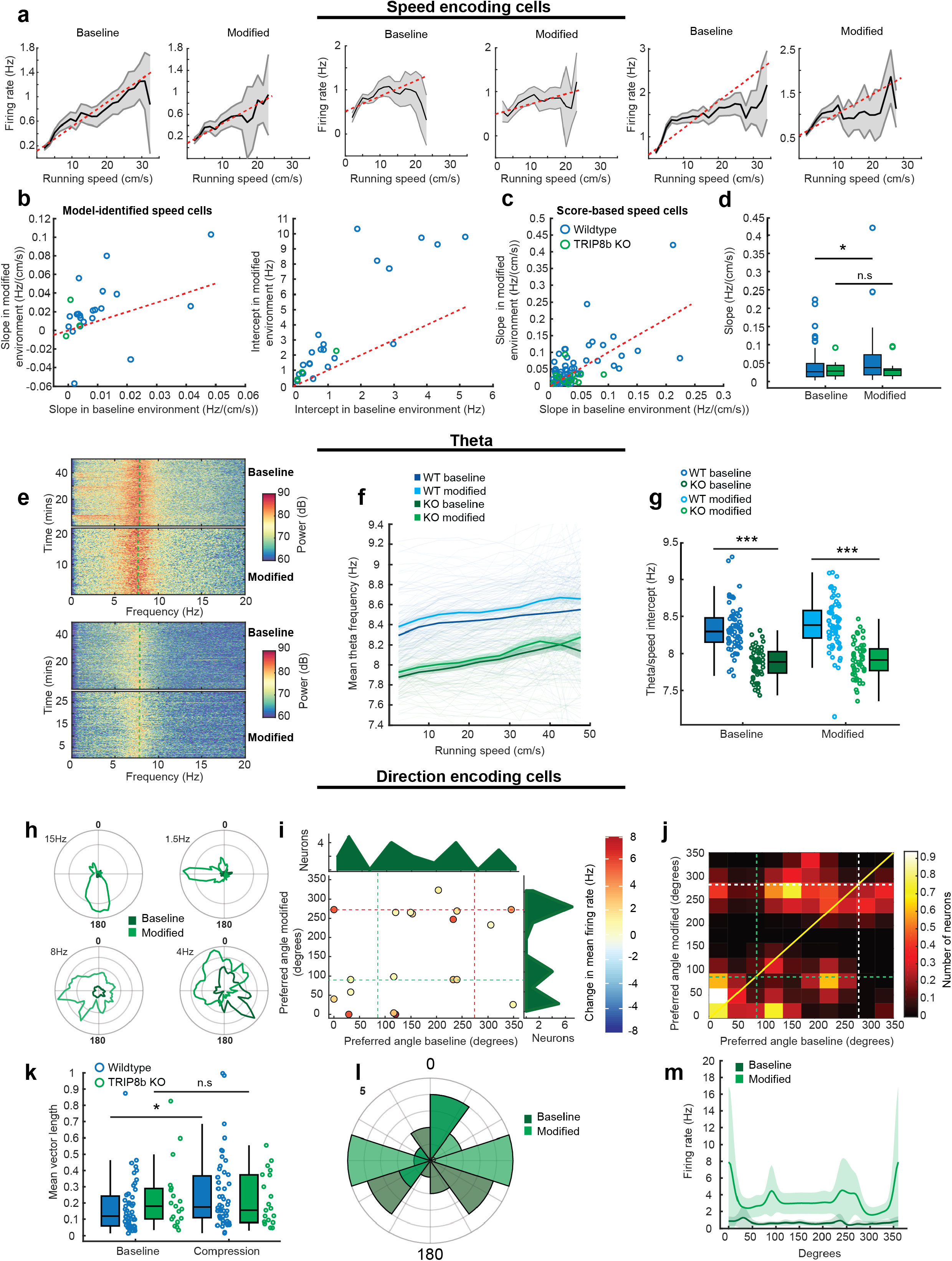
TRIP8b KO speed signals are less sensitive to environmental perturbation while directional signals remain malleable. **a,** Three example S-encoding cells in the baseline (left panels) and modified (right panels) environment for the compression condition. Dotted red line indicates the linear function fit through the data. Solid black line indicates the mean response, with gray denoting ± SEM. **b,** Scatterplot of the slope (left) and intercept (right) of KO S-encoding cells (n = 3) in the baseline and modified environment for the compression condition. S-encoding cells from the WT compression experiments shown in blue for comparison. **c,** Scatterplot of the of KO and WT speed cells defined using traditional speed scores. There was no significant difference in the slope of score-identified KO speed cells between baseline and modified environments. **d,** Boxplots of the slope of KO (green) and WT (blue) score-identified speed cells in the baseline and modified (compressed) environment. Box illustrates median ± 25^th^ and 75^th^ percentiles and whiskers 1.5x the interquartile range. Slopes beyond this range are illustrated as individual points. **e,** Example spectrograms showing LFP power in the 0-20Hz range in two sessions recorded from TRIP8b KO animals in the baseline (top panels) and modified (bottom panels) environments. The mean theta frequency over each session is shown as a superimposed dotted green line. **f,** Mean theta frequency by running speed for KO (greens) and WT (blues) animals in the baseline and modified (compressed) environments. Solid lines show the mean, shaded region is ± SEM. Individual sessions are shown as faint lines. **g,** The intercept of linear regressions of the running speed/theta frequency relationships for sessions recorded from WT animals (blues) and KO animals (greens) in the baseline and modified (compressed) environments. There was no difference in the intercept between baseline and modified environments in KO animals (KO intercept ± SEM (Hz): baseline = 7.87 ± 0.03, modified = 7.91 ± 0.03, Z = 1.72, p = 0.086). WT theta intercept was greater than KO theta intercept in both baseline (Z = 7.79, p < 0.001) and modified (Z = 7.34, p < 0.001) environments. Data are presented as jittered scatterplots and summarized as boxplots. Box illustrates median ± 25^th^ and 75^th^ percentiles and whiskers 1.5x the interquartile range. **h,** Polar plots of KO H-encoding cells in the compression condition. Top left numbers show the firing rate indicated by the outermost ring. **(I)** Scatterplot of preferred phase angle (degrees) of KO H-encoding cells between baseline and modified environments in the compression condition. The change in firing rate is indicated by colors coded for minimum (blue) and maximum (red) values. Marginal density plots (green) indicate the distribution of phase angles. The wall axes are indicated by dotted lines (green, static wall; red, moved wall). **j,** Number of KO neurons falling in each 33° directional bin in the baseline and modified environments in the compression condition. A 2-bin Gaussian filter is applied to the data. The wall axes are indicated by dotted lines (green, static wall; white, moved wall). A solid yellow line demarks the bins in which there was no change in direction. **k,** The mean vector length of H-encoding WT and KO cells in the baseline and modified environments. Data are presented as in (g), except all data are shown as jittered points adjacent to the respective boxplot. **l,** Polar histograms show the number of cells with a given preferred phase angle in baseline (dark green) and modified (light green) environments, with scale (n) indicated by the number on the top left. **m,** Mean firing rates for all H-encoding cells in the compression condition over all exposures (n = 19). Solid lines are mean firing rates, ± SEM is shown as a shaded region. The wall axes are indicated by dotted lines (green, static wall; red, moved wall). All panels: *p < 0.05, ** p < 0.01, ***p < 0.001, n.s., not significant.

A comparable lack of change in speed coding was also observed for oscillatory activity in the theta-band (5 – 11 Hz) in MEC. As expected from previous work, theta frequency was lower in KO compared to WT mice in baseline (session n = 53, mean frequency ± SEM: WT = 8.33 ± 0.037 Hz, KO = 7.87 ± 0.029 Hz, Z = 7.79, p < 0.001) (Figure 6e-g) ^28,29^. In contrast to WT animals, there was no difference in theta frequency between the baseline and modified environment (mean frequency ± SEM: baseline = 7.87 ± 0.029 Hz, modified = 7.91 ± 0.033 Hz, Z = 1.72, p = 0.086). There also was no difference in the slope of the running speed/frequency relationship in KO animals between baseline and modified environments (mean slope (Hz/(cm/s)) ± SEM: baseline = 0.0069 ± 0.0012, modified = 0.0076, ± 0.0010, Z = 0.35, p = 0.73) (Figure 6f-g). Together, these results point to coupled responses of speed and grid cells to changes in environmental structure and raise the possibility that MEC speed inputs are necessary to update grid cell estimates of position.

### Bias in head direction coding after environmental perturbation in TRIP8b mice

Unlike the relative inflexibility of the grid and speed code to environmental perturbations, head direction cells (mouse n = 6, H-encoding cell n = 19) in KO mice showed signatures of reorganization in line with those observed in WT mice (Figure 6h). There was no difference between WT and KO in the proportion of cells that retained their directional coding between baseline and modified environments (WT = 25.65%, KO = 25%, X^2^ = 0.012, p = 0.91) and, as in WT mice, the mean firing rate of HD cells significantly increased between environments (mean firing rate (Hz) ± SEM: baseline = 0.72 ± 0.14, modified = 3.18 ± 0.72, Z = 3.28, p < 0.001). Moreover, as in WT animals, there was a reorganization of preferred directions in the modified environment compared to baseline (Figure 6i-m). Unlike WT animals, while cells reorganized, there was no clear directionality in either baseline or modified environments (Hodges-Anje test: baseline, m = 6, p = 0.72; modified, m = 5, p = 0.4) (Figure 6l). These changes in the KO head direction cell population contrasts with the insensitivity of grid cells and MEC speed codes to environmental perturbation, pointing to head direction cells as contributing a velocity signal that is uncoupled from speed signals.

## Discussion

We observed changes in the MEC speed signal that closely matched the changes in grid cells after environmental deformation. Critically, when there was an asymmetric change in grid spacing, there was a matching asymmetric change in the speed cell signal. Complementary to this, expanding the environment from an asymmetric rectangle to a symmetric square caused a symmetric change in grid spacing and a matching symmetric change in the speed cell signal. These results contrast with the previous proposal that speed cell coding is context-invariant ^4^. Moreover, our results support network level attractor models of grid cell formation ^3,15,17–19^, specifically the prediction that asymmetric re-scaling of grid cells can result from asymmetric changes in velocity signals ^15^. Importantly, this change in MEC speed coding does not rule out an influence of boundary landmarks or boundary-tethered shifts in grid phase on grid re-scaling, but rather reveals that re-scaling in speed signals provides an additional, complementary mechanism for grid distortions and re-scaling ^8,9,12–14,31^.

In contrast to the changes in MEC speed cells, MEC head direction cells appeared to organize in a manner that could provide information about the stability of cues in the environment or asymmetry in environmental geometry. Moreover, the directional bias developed in the rectangular environment only after we began perturbing the environment, regardless of whether this was the familiar or novel environment, suggesting that learning is required for MEC head direction re-organization. These features of head direction re-organization support the idea that MEC head direction cells may not serve as the movement-direction signal required by grid cell models ^32^. Even so, this does not rule out that the directional component of a velocity signal may be represented by a small subset of MEC cells, or derive from head direction signals originating in another region ^26,32–35^. Finally, the bimodal nature of the head direction organization we observed raises the possibility that MEC head direction cells provide directional information that is relative to an internal reference frame ^36^. This type of code is reminiscent of head direction signals in retrosplenial cortex, a brain region with strong reciprocal MEC connections, and that contains bidirectional head direction cells that anchor their preferred firing angle to local landmarks ^37,38^.

The mechanism by which speed signals asymmetrically re-scale to reflect changes in environmental geometry remains unknown. Previous work has found that speed cells determine a component of their movement estimates from visual cues ^39,40^, which could involve optic flow information derived from movement relative to static visual landmarks or visual features on the ground plane ^41,42^. Thus, one possible mechanism for the observed speed cell asymmetry is that MEC speed cells use learned visual angles of landmarks to estimate the relative motion between the animal and environment ^41,42^. Regardless of the mechanism however, our data indicate that speed cells are not static across environmental conditions as previously suggested but rather, can re-scale in a manner similar to grid patterns to reflect changes in the geometry of the spatial environment ^4,10,11^

Finally, the loss of re-scaling in spatial and speed codes after KO of TRIP8b sheds light on the potential mechanisms underlying these phenomena. HCN channel expression controls membrane integration kinetics and thus, the loss of HCN channels could impact the sensitivity of MEC grid and speed cell responses to changes in self-motion cues ^43,44^. HCN channels also play a key role in generating rhythmic activity in the medial septum ^28,29^, which contains MEC projecting neurons that encode running speed in their firing rate ^45^. As the MEC grid network likely depends on multiple inputs to estimate a speed signal ^14^, the loss of HCN channels in the medial septum could impair speed coding at the level of both rate-coded speed inputs to MEC and the frequency modulation of theta oscillations by running speed. It also remains unknown if the loss of HCN impacts processing in higher-order visual cortices, which encode differences in movement speed and the spatial frequency of stimuli ^46–48^, as well as running speed ^49^. The loss of TRIP8b also reduces the expression of HCN1 in the retina, which may result in visual features carrying less information regarding the animal’s velocity ^50^. However, determining precisely what intrinsic or network properties result in the loss of grid and speed re-scaling in TRIP8b KO animals awaits the development of approaches that can specifically target these cell-types.

## Acknowledgements

LMG is a New York Stem Cell Foundation – Robertson Investigator. This work was supported by funding from the New York Stem Cell Foundation, NIMH MH106475, James S McDonnell Foundation and a Klingenstein-Simons award to LMG, the Philip Wrightson Postdoctoral Fellowship from the Neurological Foundation of New Zealand awarded to RGM, and an NSF Graduate Research Fellowship awarded to CM. We thank A Borrayo and A Diaz for histology assistance.

## Author Contributions

LMG and RGM conceptualized experiments and analyses. CM and RGM performed chronic implantations, and collected and analyzed in vivo data. KH provided support on analyses. DMC provided the TRIP8b KO mouse line. LMG and RGM wrote the paper with feedback from all authors.

## Methods

### Subjects

All procedures were approved by the Institutional Animal Care and use Committee at Stanford University School of Medicine. Male C57BL/6 (WT, n = 31) and TRIP8b knockout (KO) mice (n = 11; donated by D. Chetkovich, Department of Neurology, Vanderbilt University) were housed in groups of between one and five same-sex littermates. TRIP8b mice were generated from a C57BL/6 background, as previously described ^30^, and bred at Stanford University School of Medicine. After surgical implantation, mice were housed singly in transparent cages on a 12-h light/dark schedule. All experiments were conducted during the light phase. Animals were between 1 and 6 months old at time of surgery (18-30 g). Prior to surgery, animals had *ad libitum* access to food and water. After three days had elapsed post-surgery, animals were food-restricted to maintain ∼ 80% of their free-feeding weight. All animals continued to have free access to water. Fifteen WT mice were recorded in the environmental compression condition, and 16 WT mice were recorded in the environmental expansion condition. The experimenter was not blind to the condition of the mice (either genotype or recording geometry) during the experiments.

### Surgery

Animals were given prophylactic and initial anesthesia by intra-peritoneal injection of buprenorphine (0.1 mg/kg) and were then maintained under anesthesia via inhalation of a mixture of oxygen and between 0.5 – 3% Isoflurane. Mice were implanted unilaterally with a two tetrode, 8-channel Microdrive (Axona Ltd.) connected to 17 μm polyimide-coated 90% platinum 10% iridium wire tetrodes cut flat. Just prior to implantation, the tips of the electrodes were electroplated with platinum until their measured impedance ranged between 150-200 kΩ. During surgery the skull was thinned over the putative target site at between 3.2-3.3 mm from the midline and 0.5 mm posterior to lambda until the transverse sinus was visualized. Electrodes were then implanted 0.5 mm anterior to the sinus at an angle of between 2-6 degrees caudal in the sagittal plane. Six stainless steel jeweler’s screws were threaded into the skull at intervals around the incision site, and the Microdrive assembly was affixed to the skull using acrylic dental cement. One of the anterior skull screws was soldered to a ground wire, which was then connected to the Microdrive ground pin. After surgery, animals were continuously monitored until they recovered, then monitored on an hourly basis for the first six hours, and on a 12 hour basis for the first three days post-surgery. During these three post-surgery days, animals were administered Carprofen analgesia as required. After at least 3 days post-surgery, animals were habituated to the baseline recording environment that corresponded to their assigned group (i.e. a rectangular or square environment see “In vivo Single-Unit Data Collection”). Recordings began 7 days post-surgery.

### In vivo single-unit data collection

Animals in the compression condition were screened for single units in a 1 × 1 m square box, while animals in the expansion condition were screened in a 0.5 × 1 m rectangular box, created by the insertion of a removable wall 0.5 m into the 1 × 1 m environment. The environments were constructed of black polycarbonate, and each environment featured a white paper cue card affixed to the static wall on the opposite side from the movable wall. The moveable wall itself was constructed of the same material as the rest of the environment and was the same height as all the walls (0.5 m). Black curtains surrounded the perimeter of the recording environment. During recordings, mice were connected to an eight channel headstage that consisted of AC-coupled unity-gain operational amplifiers and connected through a counterbalanced cable to a DACQUSB pre-amplifier and system unit (Axona Ltd). Acquired signals were amplified between 5,000 and 20,000 times, and then band-pass filtered between 800 and 6,700 Hz. A threshold for signals was manually selected and the resultant spikes were stored at a sampling rate of 48 kHz. One of the channels was manually selected to be simultaneously used for recording EEG. This channel was digitally amplified 3,000-10,000 times and lowpass filtered at 500 Hz with a notch filter at 60 Hz. This channel was simultaneously recorded at high resolution at 48 kHz and lower resolution at 250 Hz. The EEG trace was saved continuously to disk. Animal position was stored alongside the electrophysiological recordings through means of tracking two groups of light-emitting diodes (LEDs); one “large” consisting of four individual LEDs and one “small” consisting of two individual LEDs. These LEDs were recorded by a camera mounted over the recording environment.

Each animal underwent no more than one condition (baseline and modified environments) per day before being returned to its home cage and then to the vivarium. During the recordings, animals were free to explore the environment and were encouraged to free forage with randomly scattered crumbled chocolate cereal. Animals were screened in the baseline environment for the presence of single units. All baseline recording sessions lasted at least 10 minutes and continued until the animal had covered more than 70% of the recording environment, or 60 minutes had elapsed. If 60 minutes elapsed and animals failed to cover 70% of the recording environment, they were removed from the baseline environment, returned to their home cages, and screened again the next day. If cells of interest were isolated during the baseline recording session, animals were immediately transferred to a modified environment (for WT mice in the compression experiments and TRIP8b KO mice, this was a 1 m x 0.5 m rectangle; for mice in the expansion experiments this was a 1 m x 1 m square). Animals were recorded in the modified environment for at least 10 minutes and until they had covered at least 70% of the environment, or 60 minutes had elapsed. If the animal had not covered > 70% of the environment after 60 minutes it was removed from the modified environment, returned to its home cage, and the data was not used except to determine the number of exposures each animal had to the modified environment. If no units of interest were uncovered or all data had been collected at a given recording depth, the tetrodes were advanced between 25 and 100 μm at the end of the recording in the baseline environment and the animal was returned to its home cage until the following day. The recording environments were cleaned immediately after each recording by spray-on and wipe-down application of Nature’s Miracle stain and odor eliminator (Spectrum Brands Inc.)

### Spike sorting and 2-D position estimation

Spikes were analyzed offline using a vendor-specific cluster-cutting software (TINT, Axona Ltd). Initial clustering was performed manually by examining spikes one dimension at a time in multidimensional feature space. Initial clustering was done using amplitude comparison of each channel on a given tetrode against all other channels. Spikes and positions recorded from epochs during which animals ran < 2 cm/s or > 100 cm/s were excluded. Only clusters with more than 100 spikes remaining after this exclusion were used for further analysis. Cluster quality and separation distance was calculated for each putative cell post-hoc. Cluster separation was calculated by determining the distance, in Maholonobis space, between the putative spikes belonging to one cells from spikes belonging to all other cells recorded on the same tetrode ^51^. Cluster matching between baseline and compressed environments was performed manually using cluster location and waveform comparison. The center of mass of each resulting cluster in voltage space was then determined. The center of mass of each cluster in the baseline and modified environments was used to determine drift in the center of mass between the recording sessions (See Figure S1). Putative interneurons were separated from putative excitatory cells by examining spike rate and spike width. Digitized position data was binned into 2.5 × 2.5 cm bins and smoothed using a 21-sample boxcar filter using a 400 ms window, with ten samples on each side. A 5 × 5 bin quasi-Gaussian kernel was used to filter time and spiking maps. The firing rate of each cell was calculated from these smoothed maps by dividing the number of spikes in each bin by time spent in the respective bin. The peak rate was determined from the bin with the highest firing rate. To calculate scores (i.e. grid, speed, and mean vector length) maps were adaptively smoothed ^52^.

### Histology and determination of recording positions

Data collection ceased when the signal to noise ratio precluded the successful isolation of spikes or the animal was exposed to the modified environment for more than 13 sessions - whichever came sooner. Due to the angle of implantation of the tetrodes, the development of noise on many channels typically indicated that the electrodes had passed through MEC and into the extracortical space. Mice were given an acute intraperitoneal overdose of sodium pentobarbitone and transcardially perfused with 0.9% saline to remove blood from the circulatory system, and then with 4% paraformaldehyde to fix the tissue. Animals were decapitated and left for several minutes to allow for shrinkage of the brain tissue. The tetrodes were then wound as far up their travel allowed, and then the skull was removed allowing the brain to be removed. Brains were then stored in 4% paraformaldehyde solution until sectioning. In all cases, an attempt was made to electrolytically lesion the final position of the electrodes. The lesions were made after overdose by passing 20-μA current for 10-15 seconds through two of the channels on each tetrode. In animals for which the electrodes had exited the brain, no lesion was evident, and the electrode tracks were used to determine the sites of recording.

For sectioning, brains were removed from the fixation solution, the hemisphere of interest was isolated by hemispheric transection of the brain in the sagittal plane, and the brains were rapidly frozen. Brains were sagittally sectioned into 40 μm sections using a ThermoFisher HM 550 cryostat, mounted, and stained with Cresyl violet. Mounted stained sections were then manually inspected microscopically to visualize the electrode tracks and determine the path of the electrodes. In cases where the electrode track ran to the edge of the slice, the edge of cortex where the track terminated was used as the final electrode position. In mice for which a lesion was evident, the center of the lesion was determined to be the final position of the electrodes. The border of MEC was determined from the lateral position of the electrodes which was determined primarily by the cytoarchitecture of the nearby hippocampus and cross-referenced with a stereotaxic atlas ^53^.

### Shuffling

For each cell in the dataset of 555 cells that were recorded in the baseline and modified environments, 100 shuffles of the spike trains were produced, and each shuffled spike train was used to compute a grid score, a speed score, a directional mean vector length, and a speed stability score. These variables were calculated in the same way as the original datasets. Each cell’s spike train was shuffled by transposing each spike a random interval from 20 seconds prior to its true position, or up to 20 seconds ahead of its position. From the resulting arrays of 55,500 shuffles, the 95^th^ percentile was determined using the *profile* function in the statistics toolbox of MATLAB 2017b

### Tuning curve classification and thresholding

As we limited the number of times an animal could experience the modified environment, we first screened cells in the baseline to identify cells of interest to be recorded in the modified environment. This screening process used score methods to classify functionally-defined cell types ^4,23^. Spatial maps were produced from all recordings such that the location of the animal was divided into 2 cm × 2 cm bins, and the mean spikes/bin for the recording was determined. Gridness scores were calculated from the autocorrelation of the rate map at 60 and 120 degrees compared to 30 and 90 degrees, as previously described ^23,54^. Speed scores were calculated as the Pearson correlation between the smoothed instantaneous firing rate of the cell and the running speed of the animal in 20 ms time bins, as previously described ^4^. The stability of the firing rate/running speed relationship was calculated as in Kropff, et al. ^4^; briefly, each recording session was divided into equal quarters, and the mean firing rate for each 2 cm/s speed bin from 5-50 cm/s was calculated. Stability was calculated as the mean correlation coefficient over each tuning curve. Directionality was determined from the mean vector length of a cell’s firing over all directions; more unidirectionally specific cells produced a greater mean vector length, and vice versa. Briefly, mean vector length was calculated from the mean firing rate in each 0.5 degree directional bin that was then smoothed using a 14.5 degree mean window filter as in Mallory, et al. ^22^. For the purposes of determining whether cells were of interest for inclusion in the environmental manipulation sessions, we used conservative score-based cutoffs informed from shuffled score distributions from previous similar experiments ^21,22^. Specifically, cells were classified as of interest if they had a grid score > 0.35, speed score > 0.05, or a mean vector length over 0.2.

### Cell classification using the linear-non-linear Poisson spiking (LNP) model

After data collection, we employed a statistical model based approach to identify which neurons carried significant information about position (P), head direction (H), and running speed (S) (Hardcastle et al., 2017; Mallory et al., 2018). Following previous work, we determined the minimum set of variables encoded by each neuron by using a greedy forward-search feature selection method in combination with 10-fold cross-validation (Hardcastle et al., 2017). In short, variables were added to the model if they significantly improved model-fit (quantified through log-likelihood increase from a baseline model) on held out data. Models were fit by minimizing the negative Poisson log-likelihood of the observed data (spike-trains) given the model prediction using MATLAB’s fminunc function. Models that did not significantly encode any variables (e.g. model performance was not above a baseline mean-firing rate model) were labeled “unclassified.” Regressors for each variable (e.g. position) were computed based on spline interpolation between set control points (position n = 144, head direction n = 12, speed n = 21). After the model-fitting procedure, we computed model-derived tuning curves for each encoded variable using the set of learned parameters. To distinguish different types of speed-encoding, we fit an additional model that split speed into its directional components; e.g. the model considered position (P), head direction (H), speed in the x-direction (Sx), and speed in the y-direction (Sy) as external covariates that could predict cell spiking. Model selection and model-fitting were otherwise identical.

### Analysis of grid cells

Grid scores were calculated from the rate maps of each cell. Cells were classified as grid cells if their score exceeded the 95^th^ percentile of the distribution of grid scores produced from the entire shuffled data set (55500 shuffles). We considered only those cells with a threshold-crossing grid score (P95 = 0.35) that also significantly encoded position (P) according to the LNP model. The distance from the center of each autocorrelogram to each of the surrounding six autocorrelation nodes was used to compute the grid spacing; the pair of nodes closest to the axis that was modified (i.e. compressed or expanded) was used to determine node spacing in the “modified” axis, and the set of nodes closest to the orthogonal axis were used to determine the spacing in the “static” direction (mean node orientation (degrees) ± SEM: WT compressed orientation = 57.04 ± 3.78, static orientation = 4.11 ± 5.94). For the sole purpose of fitting an ellipse, the central set of six autocorrelation nodes were manually designated (the experimenter was blind to the compression or expansion condition and genotype of the rate maps during this selection) and were used to fit an ellipse for the purpose of determining the ellipticity of the grid pattern using a least-squares model ^11,55^. The ellipticity (ε) of each cell was then computed as a ratio between the semi-major (α) and semi-minor (β) axes of this fitted ellipse.

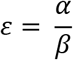

This produced an ellipticity value between 1 (a perfect circle) and infinity (^∞^, the point at which the ellipse becomes parabolic).

In order to determine the amount of deformation of the grid pattern in the modified environments compared to the baseline environments, we expanded the compression map (in the compression experiments) or compressed the expansion map (in the expansion experiments) using the *imresize* function in MATLAB 2017b. This procedure produced a stretched or compressed (rescaled) version the modified environment map over an array of values that each differed by a single spatial bin width (2.5cm). Each re-scaled map was also translated in 1 cm increments (up to 20 cm) in both the x and y dimensions. Each re-scaled and translated map was then correlated against the corresponding baseline map. From the resultant matrix of correlations (Figure S2), the combination of re-scaling and x and y translation that produced the optimum correlation (Rho) was selected to derive the proportion of deformation of the modified environment map at this point. This value (Lambda, λ) was a value ranging between 0 (best correlation when the modified map was not resized) and 1 (the best correlation was achieved when the modified map was resized such that it matched the dimensions of the corresponding baseline map).

### Analysis of direction-encoding cells

During initial screening in the baseline experiments, cells of interest were determined based on their head direction mean vector length score, as previously described. Directionality in all of the 555 cells in the dataset was determined post-hoc using the LNP model. This method captured cells that were considered traditionally “directional” using the score-based criterion (All model directional cells (n = 171): mean vector length ± SEM, baseline = 0.29 ± 0.02, modified = 0.25 ± 0.02), but also uncovered cells with unconventional tuning curves that nevertheless still encoded direction. The LNP model produced a direction-conditioned tuning curve for each cell, which was divided into 60 bins of 0.1047 radians (6 degrees). The “preferred” direction of each directional cell was therefore determined from these tuning curves for each recording as the direction of maximal firing. The mean firing rate of each neuron for each recording was determined as the average over the firing rate in each of the 60 equally-sized directional bins for each recording.

### Analysis of speed-encoding cells

During initial screening in the baseline experiments, cells of interest were determined based on their tuning curve-based speed score, as previously described. Cells that coded speed were identified post-hoc by the LNP model. As with H-encoding cells, these model-identified cells often included cells that had a super-threshold speed score, but also included some cells that would ordinarily have been overlooked by traditional shuffling methods (All S-encoding cells (n = 55) speed score ± SEM: baseline = 0.162 ± 0.018, modified = 0.105 ± 0.018) The speed-conditioned spike tuning curves of these cells was derived from the model; and provided the speed/firing rate relationship. The overall slope and intercept of this relationship was determined for each speed encoding cell by the fitting of first-degree (linear) polynomial through these data by using the *polyfit* function in MATLAB. In order to determine the speed/firing rate relationship of these cells in each individual axis, the 2D speed-conditioned tuning curve in the initial LNP model was reduced to two 1D curves, each conditioned on speed in only the x or y axis. Slope and intercept of these relationships was determined as for the overall 2D model. During recording, cells identified as interneurons were deliberately excluded from the recording sessions, so all speed cells were putative excitatory cells. We restricted our analyses to those cells that positively encoded speed in the baseline environment (that is, the slope of their firing rate/running speed relationship was positive).

### Analysis of theta-band EEG

EEG was recorded in baseline and modified environments simultaneously with single units. Sessions were included in analysis if both baseline and modified recording sessions met the same criteria as single units (≥ 70% coverage of the environment after epochs at which the movement of the animal was less than 2 cm/s or over 100 cm/s were excluded). Analyses of theta band EEG were carried out using data sampled at 250 Hz. EEG was bandpass filtered between 5 and 11 Hz and subjected to a Fourier transform in order to produce a power spectrum. The frequency at which the power peak occurred was determined for both the baseline and modified recording sessions. Instantaneous power, phase, and frequency were derived using a Hilbert transform. We determined the slope and intercept of the running speed/theta frequency relationship by linearly regressing these data using a least-squares method.

### Statistical Analyses

All the statistical tests were two-sided, with the exception of the LN model determination of variable coding, which used a one-sided rank test. All comparisons were tested for parametricity using a Lilliefors test. In all cases for which the compared data did not conform to a normal distribution, nonparametric tests were used to assess differences between groups. Wilcoxon sign-rank tests were used for paired data, and Wilcoxon rank-sum tests were used for non-paired data. When comparing differences between distributions, a two-sided Kolmorgov-Smirnov test was used. Directional data was analyzed using the *CircStat* toolbox for MATLAB ^56^. To determine whether directional data was uniformly distributed or not, either Rayleigh’s test (for non-bimodal data) or a Hodges-Anje Omnibus test was performed. Watson-Williams tests were used to determine whether directional distributions differed. While this test assumes a von Mises distribution, it has been demonstrated to be robust to departures from this assumption ^57^.

