## Supplemental Figures for "Entorhinal velocity signals reflect environmental geometry"

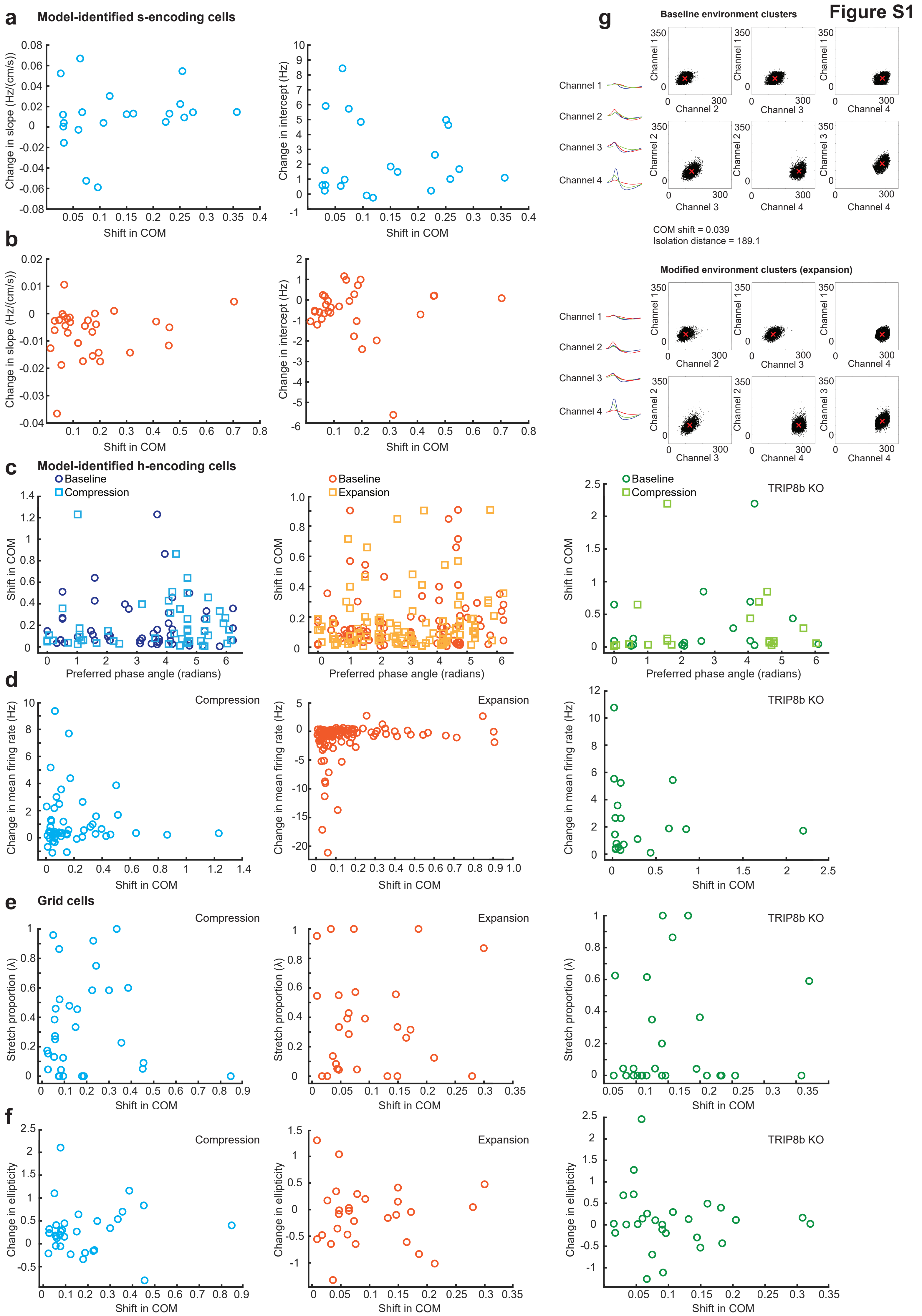
Figure S1
a
Model-identified s-encoding cells
Change in slope (Hz/(cm/s))
Change in intercept (Hz)
Shift in COM
b
c
Model-identified h-encoding cells
Baseline
Compression
Expansion
TRIP8b KO
Shift in COM
Preferred phase angle (radians)
d
Change in mean firing rate (Hz)
e
Grid cells
Stretch proportion (λ)
f
Change in ellipticity
g
Baseline environment clusters
Channel 1
Channel 2
Channel 3
Channel 4
COM shift = 0.039
Isolation distance = 189.1
Modified environment clusters (expansion)

**Figure S2**

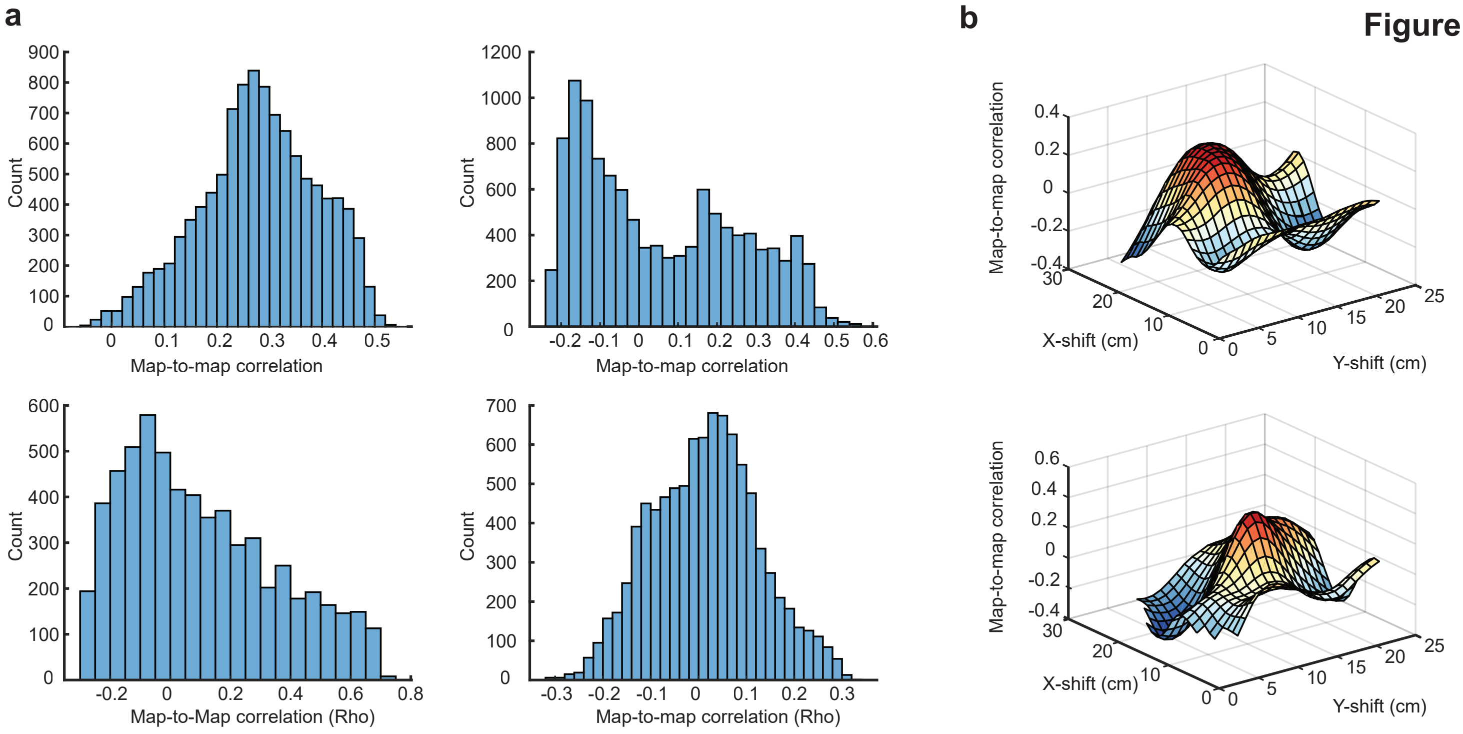
a
b
Count
Map-to-map correlation
Map-to-Map correlation (Rho)
Map-to-map correlation (Rho)
Map-to-map correlation
X-shift (cm)
Y-shift (cm)

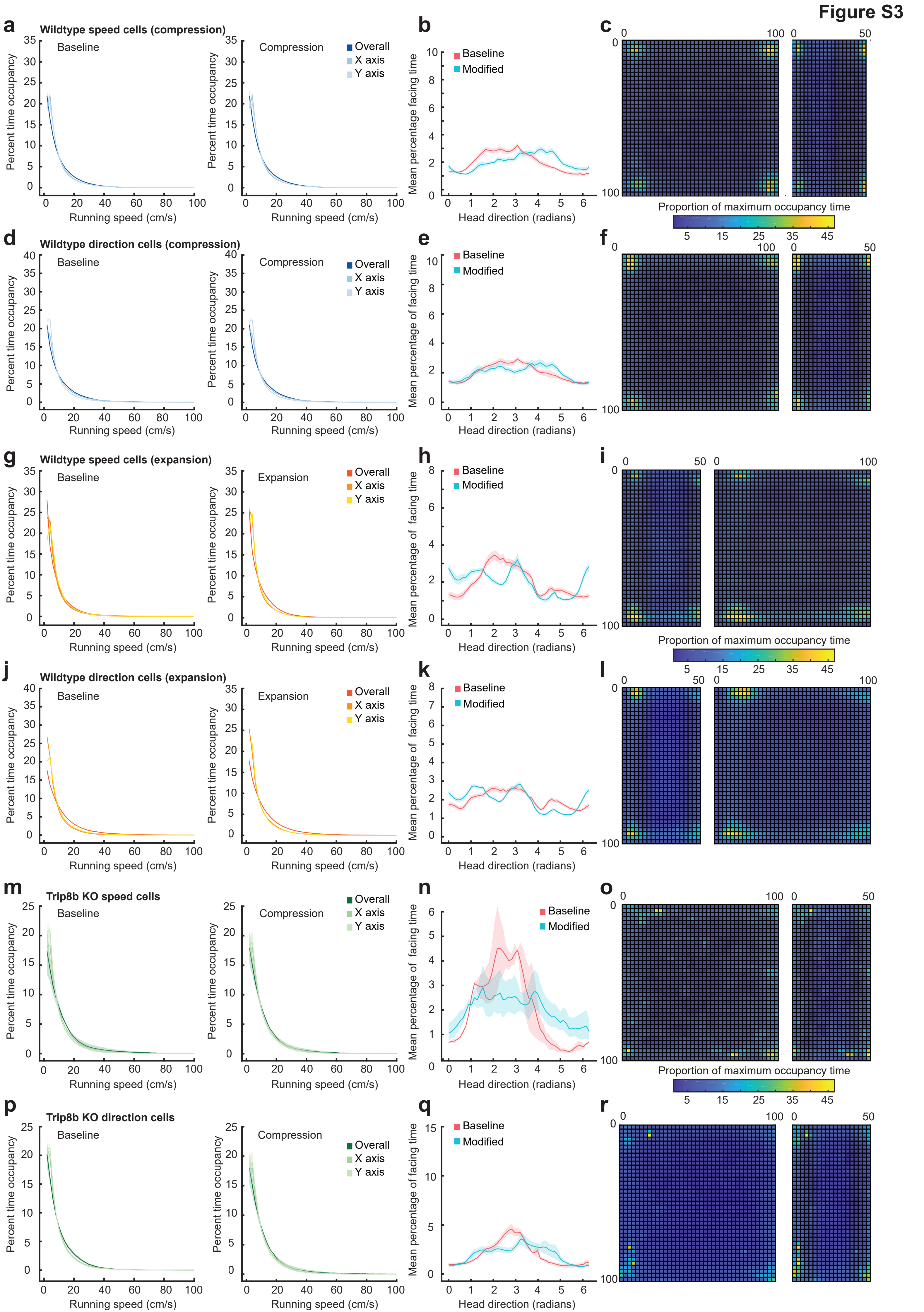
Figure S3
a
Wildtype speed cells (compression)
Baseline
Compression
Overall
X axis
Y axis
Percent time occupancy
Running speed (cm/s)
b
Baseline
Modified
Mean percentage facing time
Head direction (radians)
c
Proportion of maximum occupancy time
d
Wildtype direction cells (compression)
e
Mean percentage of facing time
f
g
Wildtype speed cells (expansion)
Expansion
h
i
j
Wildtype direction cells (expansion)
k
l
m
Trip8b KO speed cells
n
o
p
Trip8b KO direction cells
q
r

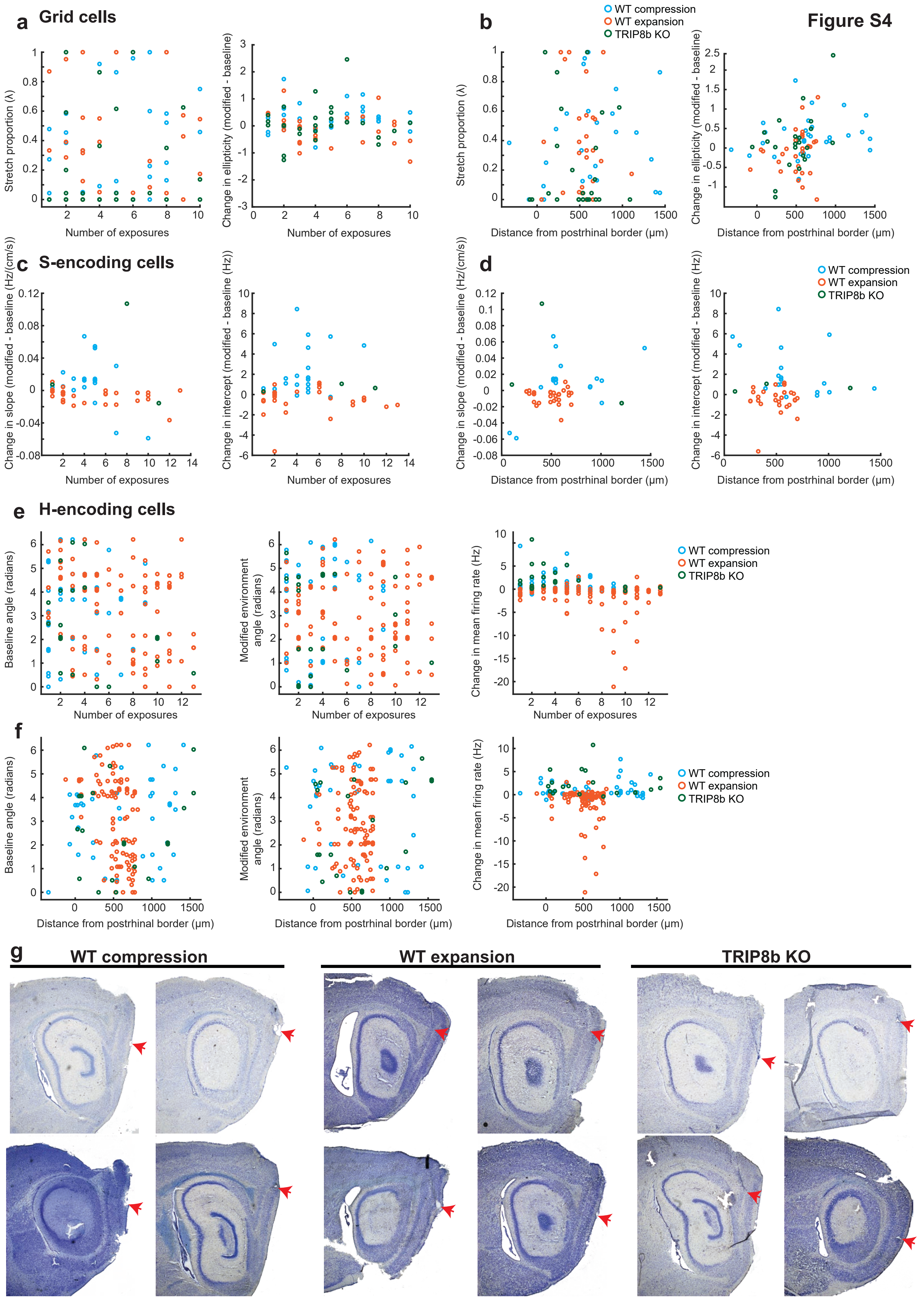

Figure S4
a Grid cells
Stretch proportion (λ)
Change in ellipticity (modified - baseline)
Number of exposures
b
WT compression
WT expansion
TRIP8b KO
Distance from postrhinal border (μm)
c S-encoding cells
Change in slope (modified - baseline (Hz/(cm/s)))
Change in intercept (modified - baseline (Hz))
d
e H-encoding cells
Baseline angle (radians)
Modified environment angle (radians)
Change in mean firing rate (Hz)
f
g
WT compression
WT expansion
TRIP8b KO

Figure S5

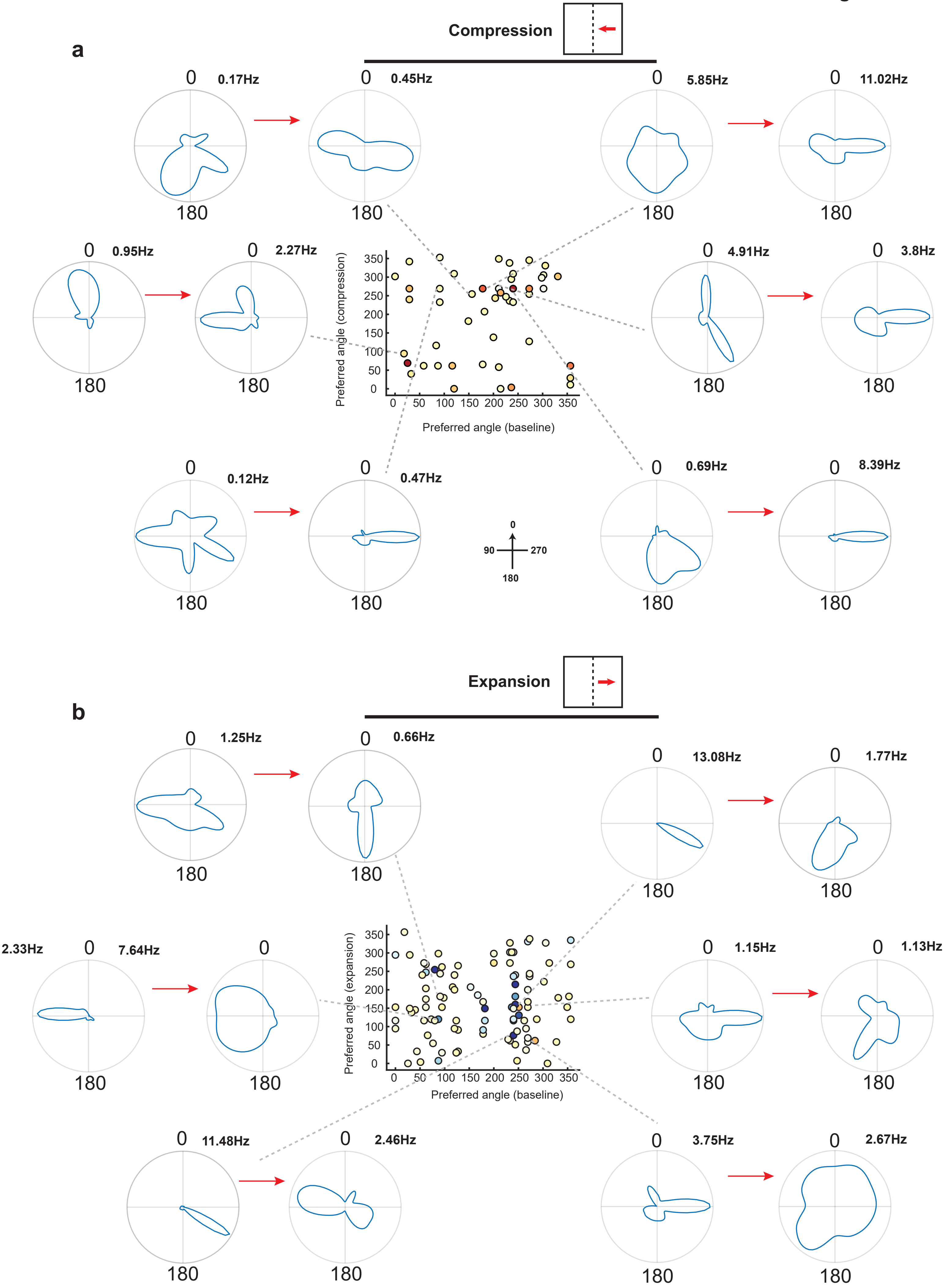

a
Compression
0.17Hz
0.45Hz
5.85Hz
11.02Hz
0.95Hz
2.27Hz
4.91Hz
3.8Hz
0.12Hz
0.47Hz
0.69Hz
8.39Hz
Preferred angle (compression)
Preferred angle (baseline)
0
90
270
180
b
Expansion
1.25Hz
0.66Hz
13.08Hz
1.77Hz
2.33Hz
7.64Hz
1.15Hz
1.13Hz
11.48Hz
2.46Hz
3.75Hz
2.67Hz
Preferred angle (expansion)
Preferred angle (baseline)

**Figure S6**

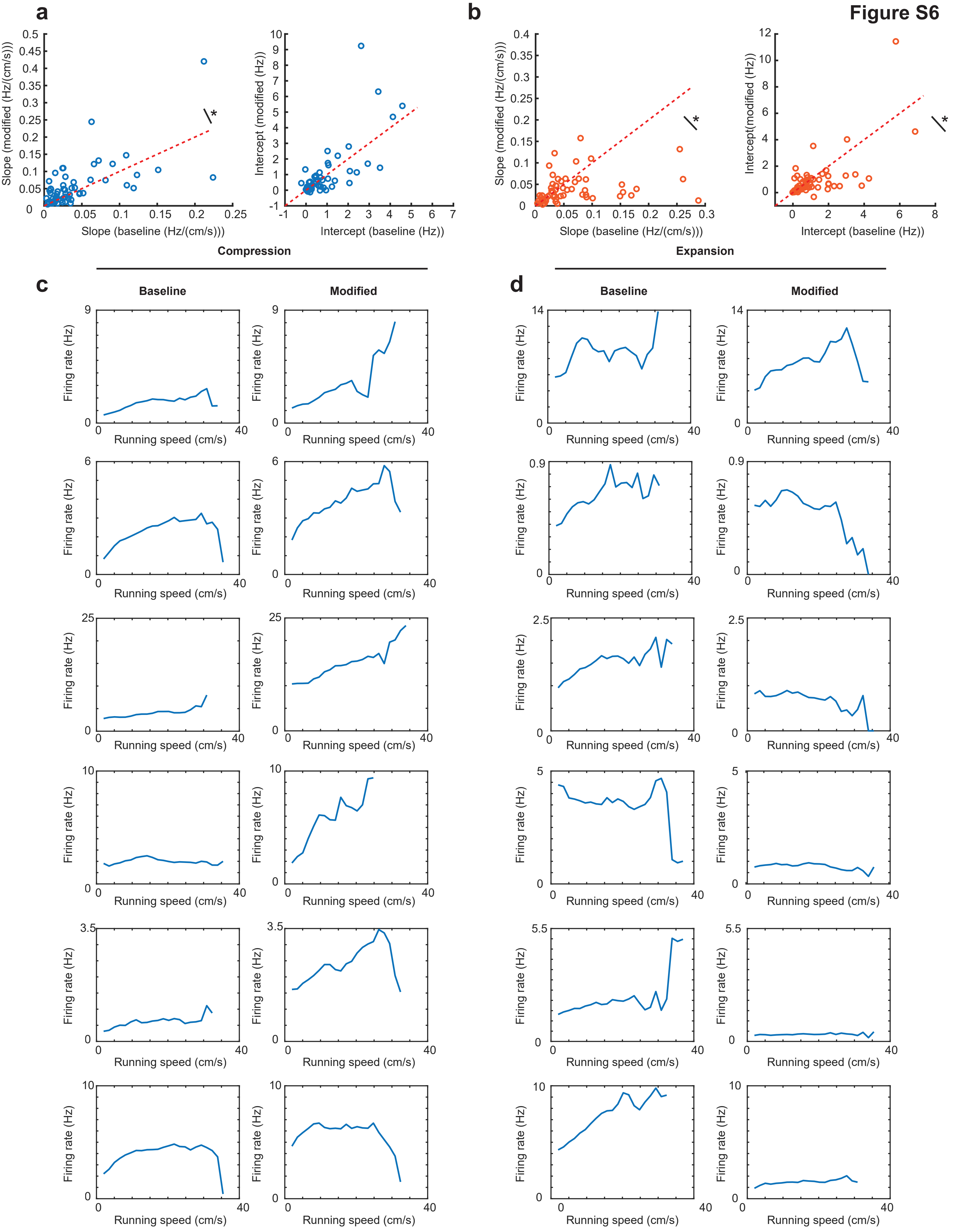
a
b
Slope (modified (Hz/(cm/s)))
Slope (baseline (Hz/(cm/s)))
Intercept (modified (Hz))
Intercept (baseline (Hz))
Intercept(modified (Hz))
*
Compression
Expansion
c
d
Baseline
Modified
Firing rate (Hz)
Running speed (cm/s)

### Supplemental Figure Legends

**Figure S1. Cluster center of mass shift was unrelated to experimental findings; Related to Figures 1, 2, 3, 5, and 6. a,** Shift in the center of mass (COM) of spikes of clusters belonging to S-encoding cells in the compression experiments against the change in slope of the speed/firing rate relationship (left panel) and the change in the intercept (right panel) between baseline and modified environments. There was no relationship between COM shift and slope change (r = 0.18, p = 0.44) or COM shift and intercept change (r = -0.05, p = 0.83). **b,** As in (a), but for S-encoding cells recorded during the environmental expansion experiments. As in (a), there was no relationship between COM shift and slope change (r = 0.14, p = 0.44) or COM shift and intercept change (r = -0.04, p = 0.82). **c,** Shift in COM against the preferred phase angle of H-encoding cells in both the baseline (circles) and modified (squares) environments. There was no relationship between preferred angle and COM shift for WT compression cells (left panel; baseline r = -0.02, p = 0.91, modified r = 0.08, p = 0.57); WT expansion cells (middle panel; baseline r = 0.06, p = 0.54, modified r = 0.13, p = 0.20); or TRIP8b KO cells (right panel; baseline r = 0.17, p = 0.48, modified r = 0.03, p = 0.90). **d,** Shift in cluster COM against the change in mean firing rates of direction encoding cells between baseline and modified environments. There was no relationship between COM shift and firing rate change in the WT compression experiments (left panel, r = -0.08, p = 0.60) or TRIP8b KO compression experiments (right panel, r = -0.09, p = 0.72). There was a small association between change in firing rate and COM shift in the expansion experiments (middle panel, r = 0.20, p = 0.05), but cells with the largest change in firing rate tended to have the smallest shifts in COM. **e,** Shift in cluster COM and the stretch proportion (λ) of grid cells in the WT compression (left), WT expansion (middle) and TRIP8b KO (right panel) experiments. There was no association between COM and λ for any group (WT compression r = -0.02, p = 0.91; WT expansion r = 0.02, p = 0.91; TRIP8b KO r = 0.06, p = 0.74). **f,** Shift in COM and the change in grid ellipticity between baseline and modified environments of grid cells in the

WT compression (left), WT expansion (middle) and TRIP8b KO (right) experiments. There was no association between COM and ellipticity for any group (WT compression r = 0.17, p = 0.37; WT expansion r = -0.03, p = 0.89; TRIP8b KO r = -0.11, p = 0.57). **g,** Example cluster of a cell recorded in the baseline (top) and then modified (bottom) environments. The position of the cluster is shown on each pair of channels on a single tetrode. COM of the cluster is illustrated with a red cross. The waveform of the cell on each of the four channels is shown in blue to the left of each panel of channels. Two simultaneously recorded units are also shown; one in green and one in red.

**Figure S2. Stretched and translated Grid Cell comparisons settle on a distinct optimum. Related to Figure 1. a,** Number of comparisons (from 8,000 combinations of stretch, x-, and y-translation) that produce a particular map-to-map correlation (rho) for four individual grid cells. In all cases, most comparisons lead to relatively poor map-to-map correlation, while the peak correlation is achieved by very few solutions. **b,** Manifold of correlations caused by all x- and y-shift combinations at one modified map stretch (λ) for two individual grid cells. As expected, the correlation between maps varies smoothly, with a defined optimal region of x- and y- shift at this λ value.

**Figure S3. Behavioral sampling: Related to Figure 1, 2, 3, 4, 5, 6, 7. a,** Mean (solid lines) ± SEM (shaded regions) percentage of time mice spent in each running speed bin for sessions during which S-encoding cells were recorded in the compression experiments. Overall speed and axis-specific speed are shown in the baseline (left) and modified (right) environments. **b,** Mean (solid lines) ± SEM (shaded regions) percentage of time mice spent facing a particular direction during sessions from which S-encoding cells were recorded during the compression experiments. Facing directions in the baseline condition are shown in red, while facing directions the modified environment are shown in blue. **c,** Mean proportion of the maximum occupancy time animals spent occupying each 2.5 cm spatial bin during sessions from which S-encoding cells were recorded. Recordings are shown for baseline square (left) and modified rectangle (right). Color coded for minimum (blue) and maximum (yellow) values. **d,e,f,** As in a,b and c, but for sessions during which H-encoding cells were recorded. **g,h,I,** As in a, b and c, but for S-encoding cells recorded during the environmental expansion experiments. In this case the baseline rectangle is shown on the left panel of (I), while the modified square is shown on the right panel. **j,k,l,** As in g,h, and i, but for sessions during which H-encoding cells were recorded. **m,n,o,** As in a,b, and c, but for sessions during which S-encoding cells were recorded from TRIP8b KO animals. Note the more variable direction sampling due to the lower number of sessions (4). In (o) the baseline square is shown on the left, while the modified rectangle is shown on the right. **p,q,r,** As in m,n and o, but for sessions during which H-encoding cells were recorded.

**Figure S4. Main experimental effects compared by number of exposures to the modified environment and recording depth. Examples of histology. Related to Figures 1, 2 , 4, 5 and 6. a,** The stretch proportion at best correlation of (λ, left) and grid ellipticity (right) of grid cells are unrelated to the number of exposures to the modified environment in the WT compression (blue) (λ, r = 0.24, p = 0.19; ellipticity, r = 0.06, p = 0.74), the WT expansion (red) (λ, r = -0.06, p = 0.78; ellipticity, r = -0.27, p = 0.17), and TRIP8b KO compression (green) (λ, r = 0.06, p = 0.78; ellipticity, r = 0.07, p = 0.72). **b,**The stretch proportion at best correlation of (λ, left) and grid ellipticity (right) of grid cells are unrelated to the distance from the postrhinal border from which they were recorded in the WT compression (λ, r = 0.13, p = 0.50; ellipticity, r = 0.19, p = 0.34), the WT expansion (λ, r = -0.05, p = 0.78; ellipticity, r =-0.04, p = 0.83), and TRIP8b KO groups (λ, r = -0.06, p = 0.76; ellipticity, r = 0.27, p = 0.19). **c,** The change in the slope (left panel) and intercept (right panel) of the firing rate/running speed relationship of S-encoding cells between baseline and modified environments is unrelated to the number of prior exposures to the modified environment in the WT compression (slope, r = -0.37, p = 0.10; intercept, r = 0.27, p = 0.22), in the WT expansion, number of exposures was unrelated to the change in slope and the change in intercept (slope, r = -0.33, p = 0.08; intercept, r = -0.01, p = 0.95). In theTRIP8b KO group, change in slope and intercept were unrelated to number of exposures to the modified environment (slope, r = -0.15, p = 0.85; intercept, r = 0.54, p = 0.46). **d,** The change in the slope (left panel) and intercept (right panel) of the firing rate/running speed relationship of S-encoding cells between baseline and modified environments is unrelated to the distance from the postrhinal border at which the cells were recorded in the WT compression (slope, r = 0.46, p = 0.06; intercept, r = -0.39, p = 0.11), WT expansion (slope, r = -0.03, p = 0.84; intercept, r = 0.08, p = 0.68), and TRIP8b KO compression (slope, r = -0.55, p = 0.45; intercept, r = 0.14, p = 0.86). **e,** The preferred phase angle of H-encoding cells in the baseline environment (left panel) was unrelated to the number of exposures to the modified environment in the WT compression (r = -0.12, p = 0.40) and TRIP8b KO compression (r = -0.003, p = 0.99) but baseline phase angles of cells in the WT expansion were negatively related to the number of exposures (r = 0.27, p = 0.005), Consistent with the observed organization of phase angles in the baseline environment. There was no relationship between modified environment phase angle and number of exposures to the modified environment (middle panel, WT compression r = 0.20, p = 0.20; WT expansion r = 0.11, p = 0.25; TRIP8b KO r = 0.15, p = 0.53). Similarly, the change in firing rate between environments was unrelated to the number of exposures to the novel environment (right panel, WT compression, r = -0.03, p = 0.84; WT expansion, r =-0.18, p = 0.07; TRIP8b KO, r = -0.31, p = 0.20). **f,** As in (e), there was no association between the distance from the postrhinal border at which cells were recorded and the preferred angle in the baseline environment (left panel, WT compression r = 0.13, p = 0.42; WT expansion r = -0.16, p = 0.10; TRIP8b KO r = 0.03, p = 0.9), preferred angle in the modified environment (middle panel, WT compression r = -0.05, p = 0.75; WT expansion r = 0.002, p = 0.98; TRIP8b KO, r = 0.30, p 0.21), or change in firing rate between environments (right panel, WT compression r = 0.15, p = 0.34; WT expansion r = -0.05, p = 0.60; TRIP8b KO r = -0.08, p = 0.74). For panels (a-f) variable *n* results from the inability to determine the position of some electrodes. **g,** Sagittal sections of mouse brain showing the electrode track and final position of electrodes for four animals in the WT compression (left panels), WT expansion (middle panels) and TRIP8b KO groups (right panels). The final position of the electrodes is marked with a red arrow.

**Figure S5. Further examples of direction-encoding cells. Related to Figure 3. a,** Six example direction-encoding cells recorded in the WT compression experiments. The model-derived tuning curve of each cell in the baseline environment is shown on the left of the pair, while the curve in the modified environment is shown on the right. Inset scatter plots are a reproduction of Figure 3B. The cell shown in the tuning curve pairs is indicated with a dashed grey line. The mean firing rate of each cell is shown at the top right of each curve plot. **b,** As in a, but for direction-encoding cells recorded during the WT expansion experiments.

**Figure S6. Examples of speed-encoding cells. Related to Figure 2. a,** Score-based speed cells; cells determined to be speed-encoding by traditional score-based methods. The slope of the firing rate/running speed relationship for WT compression cells in the baseline and modified environment (left) and intercept of this relationship in the baseline and modified environments (right). Unity in the variables in both environments is illustrated with a dotted red line. Slope (left panel) was significantly greater in the modified environment than baseline, in line with findings using a model-based approach (speed cell n = 58; slope (Hz/(cm/s)) ± SEM: baseline = 0.042 ± 0.006, modified = 0.057 ± 0.009, Z = 2.31, p = 0.02). There was no difference in intercept between environments (right panel; intercept (Hz) ± SEM baseline = 0.88 ± 0.14, modified = 0.99 ± 0.22). **b,** As in (a), but for cells identified as speed cells using scores in the expansion experiments. Slope (left panel) was significantly smaller in the modified environment compared to baseline (speed cell n = 57, baseline = 0.06 ± 0.009 Hz/(cm/s), modified = 0.04 ± 0.004 Hz/(cm/s), Z = 2.24, p = 0.025). The intercept of speed cells was also lower in the expansion compared to baseline (baseline = 1.23 ± 0.19 Hz, modified = 0.99 ± 0.22 Hz, Z = 2.1, p = 0.038). **c,** Six example raw tuning curves of speed-encoding cells in the baseline (left) and modified (right) environments in the WT compression experiments. Solid line Illustrates the mean firing rate. **d,** As in (c), but for speed-encoding cells in the WT expansion experiments.
